# The importance of species interactions in spatially explicit eco-evolutionary community dynamics under climate change

**DOI:** 10.1101/2020.03.23.003335

**Authors:** Anna Åkesson, Alva Curtsdotter, Anna Eklöf, Bo Ebenman, Jon Norberg, György Barabás

**Affiliations:** Linköping University; University of New England; Stockholm University; Linköping University; MTA-ELTE Theoretical Biology and Evolutionary Ecology Research Group

## Abstract

Eco-evolutionary dynamics are essential in shaping the biological response of communities to ongoing climate change. Here we develop a spatially explicit eco-evolutionary framework which integrates evolution, dispersal, and species interactions within and between trophic levels. This allows us to analyze how these processes interact to shape species- and community-level dynamics under climate change. Additionally, we incorporate the heretofore unexplored feature that species interactions themselves might change due to increasing temperatures and affect the impact of climate change on ecological communities. The new modeling framework captures previously reported ecological responses to climate change, and also reveals two new key results. First, interactions between trophic levels as well as temperature-dependent competition within a trophic level mitigate the negative impact of climate change on global biodiversity, emphasizing the importance of understanding biotic interactions in shaping climate change impact. Second, using a trait-based perspective, we found a strong negative relationship between the within-community variation in preferred temperatures and the capacity to respond to climate change. Communities resulting from different ecological interaction structures form distinct clusters along this relationship, but varying species’ abilities to disperse and adapt to new temperatures leave it unaffected.

## Introduction

Changing climatic conditions influence species’ ecology, such as demography, biotic interactions, and movement, as well as species’ evolutionary rates. Despite the acknowledgement of the highly important interplay between ecological and evolutionary processes for determining species distributions and survival under altered climatic conditions ^1–4^, few studies account for their combined effects ^5^. Two of the few studies addressing both types of dynamics show surprising results: inclusion of evolution potentially result in increased extinction rates when combined with dispersal ^6^, and high dispersal rates do not reduce extinctions since colonization often comes at expense of other species ^7^. Moreover, species interactions can both alleviate and aggravate the impact of climate change on species ^8^, and interact with other eco-evolutionary processes. For example, species interactions can affect a species’ evolutionary response to altered environmental conditions ^9^; and dispersal may release a species from negative interactions through migration ^10^ or increase them through invasion ^11^.

To understand species response to climate change, we need a framework that includes both ecological and evolutionary processes not only over time but also in space, while also allowing for multispecies interactions ^2^. Earlier studies strove to include a variety of relevant biological mechanisms when predicting species response to increased temperatures ^6,7^. Norberg et al. ^7^ take most of the aforementioned aspects into consideration, but at the expense of a highly simplified representation of species interactions. Thompson and Fronhofer ^6^ present a promising individual-based modeling framework, including evolution and dispersal of interacting species under climate change. Their model is in principle extensible to handle complex interspecific interactions across multiple trophic levels. However it is unclear how cumbersome their model would then become due to its computationally expensive, individual-based approach.

Here we develop a framework bringing together dispersal, adaptation and importantly, species interactions both within and between trophic levels. Furthermore, we include how species interactions themselves change because of increasing temperatures, something that has so far not been considered in an eco-evolutionary setting ^12–14^. To make the model easy to interpret and computationally efficient without having to sacrifice important details of biological processes, evolution is based on quantitative genetics ^15–19^ and space is partitioned into discrete locations.

Using this framework, we explore the effects of species interactions, dispersal, and evolutionary rates on the dynamics of species’ ranges, the overall coexistence of species, regional and global species turnover and losses, as well as the general community-wide capacity to respond to climatic change. Doing so, we find that the influence from consumers and temperature-dependent competition alter species response to increased temperatures resulting in species coexisting to a higher degree, with lower levels of species turnover and global losses being less severe. Furthermore, we demonstrate that community-wide dispersion of species’ temperature optima is a strong predictor of a community’s capacity to respond to climate change, which has implications on future management guidelines.

## Modeling framework

We consider *S* species distributed in *L* distinct habitat patches. The patches form a linear latitudinal chain going around the globe, with dispersal between adjacent patches (Figure 1). The state variables are species’ local densities and local temperature optima (the temperature at which species achieve maximum intrinsic population growth). This temperature optimum is a trait whose evolution is governed by quantitative genetics ^15–19^: each species, in every patch, has a normally distributed temperature optimum with given mean and variance. This variance is the sum of a genetic and an environmental contribution. The genetic component is given via the infinitesimal model ^20,21^, whereby a very large number of loci each contribute a small additive effect to the trait. This ensures that the total phenotypic variance is unchanged by selection, with only the mean being affected ^22^. Consequently, each species has a fixed phenotypic variance which is the same across all patches, but the mean temperature optimum may evolve locally and can therefore differ across patches (Figure 1).

**Figure 1:**
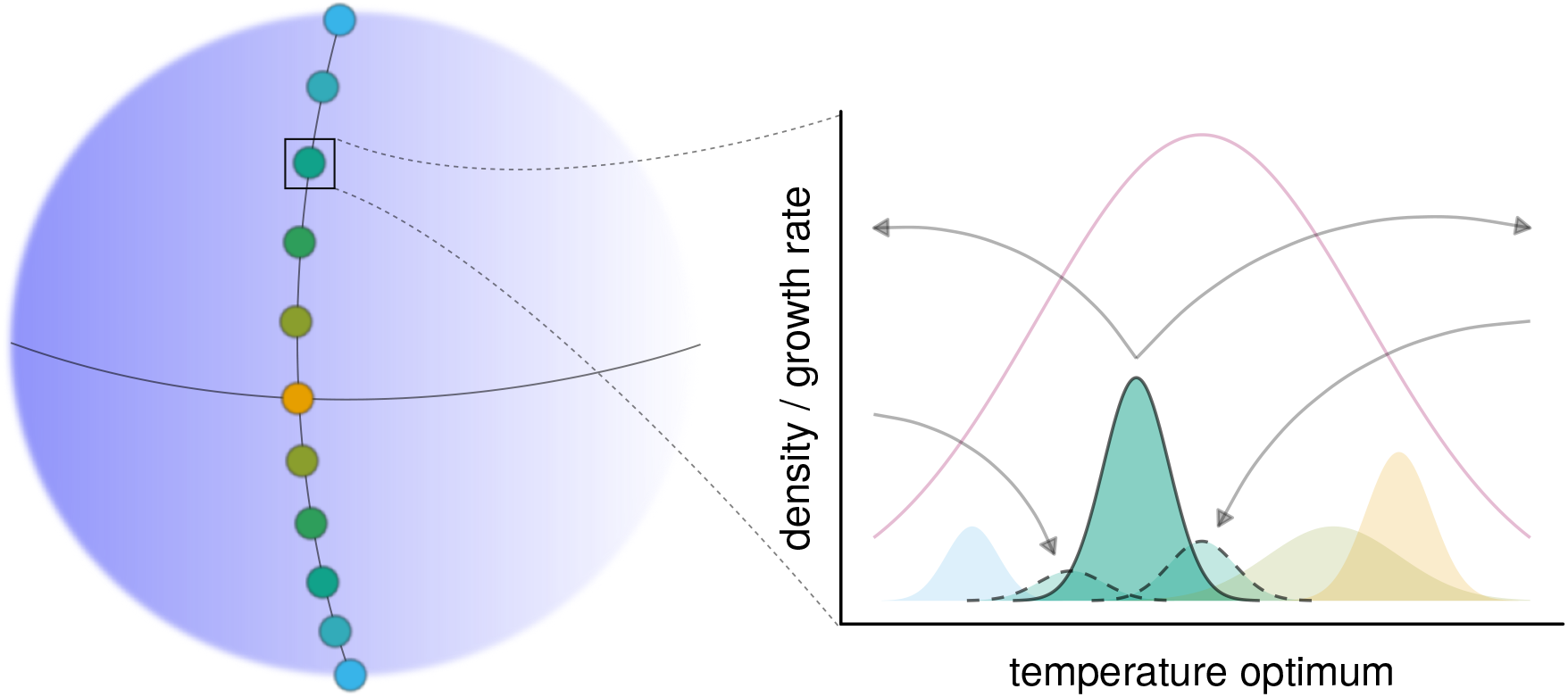
Illustration of our modeling framework. There are several patches hosting local communities, arranged linearly along a latitudinal gradient. Patch color represents the local average temperature, with warmer colors corresponding to higher temperatures. There is dispersal between adjacent patches. The graph depicts the community of a single patch, with four species present. They are represented by the colored areas showing the distributions of their temperature optima, with the area under each curve equal to the population density of the corresponding species. The green species is highlighted for purposes of illustration. Each species has migrants to adjacent patches, as well as immigrants from them (arrows from and to the green species; the distributions with dashed lines show the trait distributions of the green species’ immigrant individuals). The purple line is the intrinsic growth rate of a phenotype in the patch, as a function of its local temperature optimum. (A phenotype achieves maximum growth when its temperature optimum coincides with the temperature in the patch; similarly, a species achieves maximum growth when the mean of its temperature optimum distribution matches the local temperature.) Local population densities are changed by temperature-dependent intrinsic growth, competition with other species in the same patch, immigration to or emigration from neighboring patches, and (in certain realizations of the model) pressure from consumer species. The means of species’ trait distributions also change. First, within each patch and for each population, there is selection pressure on the mean temperature optimum to match local temperatures. Second, trait differences in the same species between patches means that immigrants (distributions with dashed lines) exert an influence on the local trait distribution after random mating. Third, in some model setups, competition between co-occurring species weakens with greater distance between them along the temperature optimum trait axis. Due to these local selection pressures, local adaptation of a species’ temperature optimum distribution may occur and, hence, differ between patches.

Within each patch, local dynamics are governed by the following five processes: 1) migration to and from adjacent patches (changing both local population densities and also local adaptedness, due to the mixing of immigrant individuals with local ones); 2) temperature-dependent intrinsic growth depending on how well species’ temperature optima match local temperatures (Figure 2**A**); 3) local competition within and between species; 4) growth from consumption; and 5) loss due to being consumed. Metabolic loss and mortality for consumers always result in negative intrinsic rates of increase, which must be compensated by sufficient consumption to maintain their populations. Each consumer has feeding links to half of the resource species (pending their presence in patches where the consumer is also present), which are randomly determined but always include the one resource which matches the consumer’s initial mean temperature optimum. Feeding rates follow a Holling type II functional response.

**Figure 2:**
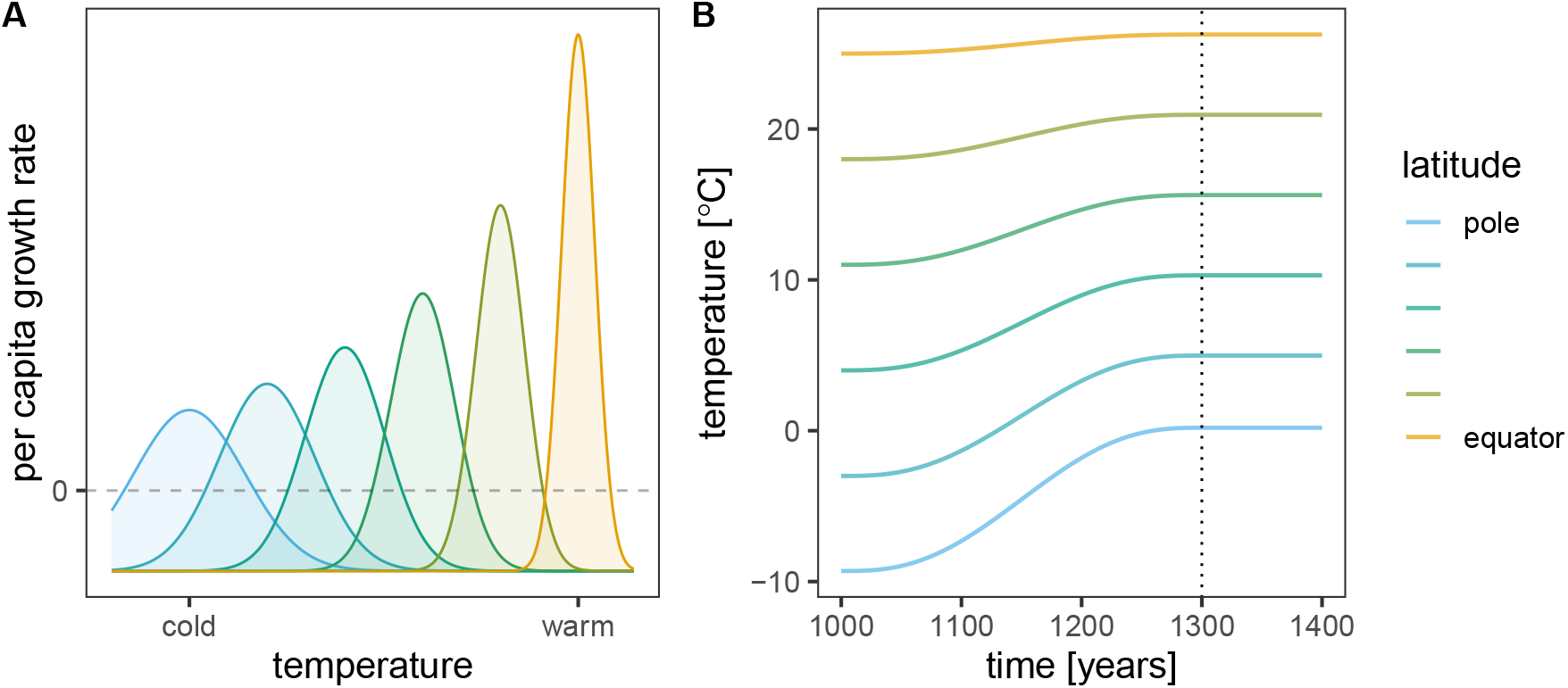
Temperature optima and climate curves. **A**: Different growth rates at various temperatures. Colors show species with different mean temperature optima, with warmer colors corresponding to more warm-adapted species. The curves show the maximum growth rate achieved when a phenotype matches the local temperature, and how growth rate decreases with an increased mismatch between a phenotype and local temperature, for each species. The dashed line shows zero growth: below this point, the given phenotype of a species mismatches the local temperature to the extent that it is too maladapted to be able to grow. Note the tradeoff between the width and height of the growth curves, with more warm-tolerant species having larger maximum growth at the cost of being viable for only a narrower range of temperatures^23^. **B**: Temperature change over time. After an initial establishment phase of 1000 years during which the pre-climate change community dynamics stabilize, temperatures increase for 300 years (vertical dotted line, indicating the end of climate change). Colors show temperature change at different locations along the spatial gradient, with warmer colors indicating lower latitudes. The magnitude and latitudinal dependence of the temperature change are based on region-specific predictions by 2100, in combination with estimates giving approximate increase by 2300, for the IPCC intermediate emission scenario^24^.

Communities are initiated with 50 species per trophic level, subdividing the latitudinal gradient into 50 distinct patches going from pole to equator (results are qualitatively unchanged by increasing the number of species; see Supplementary Information [SI], Section 5.3). We assume that climate is symmetric around the equator; thus, only the pole-to-equator region needs to be modeled explicitly (SI, Section 3.5). Temperature increase is based on predictions from the IPCC intermediate emission scenario ^24^ and corresponds to predictions for the north pole to the equator. Species are initially equally spaced, with one species per trophic level per patch, and adapted to the centers of their ranges. We then integrate the model for 3500 years, with three main phases: 1) an establishment period from 0 to 1000 years, during which local temperatures are constant; 2) climate change, between 1000 and 1300 years, during which local temperatures increase in a latitude-specific way (Figure 2**B**); and 3) the post-climate change period from 1300 to 3500 years, where temperatures remain constant again at their elevated values.

To explore the influence and importance of dispersal, evolution, and interspecific interactions, we considered the fully factorial combination of high and low average dispersal rates, high and low average available genetic variance (determining the speed and extent of species’ evolutionary responses), and four different ecological models. These were: 1) the baseline model with a single trophic level and constant, patch- and temperature-independent competition between species; 2) two trophic levels and constant competition; 3) single trophic level with temperature-dependent competition (changing based on species’ temperature optima); and 4) two trophic levels as well as temperature-dependent competition. This gives 2 × 2 × 4 = 16 scenarios. For each of them, some parameters (competition coefficients, tradeoff parameters, genetic variances, dispersal rates, consumer attack rates, and handling times; SI, Section 6) were randomly drawn from pre-specified distributions. We therefore obtained 10 replicates for each of the 16 scenarios. While replicates varied in the precise identity of the species which survived or went extinct, they turned out to vary little in the overall patterns they produced, so this was sufficient to accurately capture the model’s behavior for every scenario.

We use the results from these numerical experiments to explore patterns of 1) species range breadths, 2) global and 3) local species diversity, and 4) changes in community composition induced by climate change. In addition, we also calculated 5) the interspecific community-wide trait lag (the difference between the community’s density-weighted mean temperature optima and the current temperature) as a function of the community-wide weighted trait dispersion (centralized variance in species’ density-weighted mean temperature optima; see Methods). The *response capacity* is the ability of the biotic community to close this trait lag over time ^25^ (SI, Section 4). Integrating trait lag through time ^26^ gives an overall measure of different communities’ ability to cope with changing climate over this time period; furthermore, this measure is comparable across communities. The integrated trait lag summarizes, in a single functional metric, the performance and adaptability of a community over space and time. The reason it is related to performance is that species which on average live more often under temperatures closer to their optima (creating lower trait lags) will perform better than species whose temperature optima are far off from local conditions in space and/or time. Thus, a lower trait lag (higher response capacity) may also be related to other ecosystem functions, such as better carbon uptake which in turn has the potential to feed back to global temperatures ^27^.

## Results

We use our framework to explore how species interactions, dispersal, and available genetic variance affect species’ spatial distributions and persistence. Additionally, we investigate how these processes affect the general community-wide capacity to respond to climate change. For simplicity, we focus on the dynamics of resource species, which are present in all model scenarios. Results for consumers, in the cases they are present, are shown in the SI (Section 5.2). The model predicts large global biodiversity losses for all scenarios (Figure 3), with losses continuing even during the post-climate change period with stable temperatures, indicating a substantial extinction debt. Trophic interactions and temperature-dependent competition, respectively, reduce the number of global losses compared to the baseline model. In conjunction, the two mechanisms do lead to fewer losses than with each operating alone, but only marginally so.

**Figure 3:**
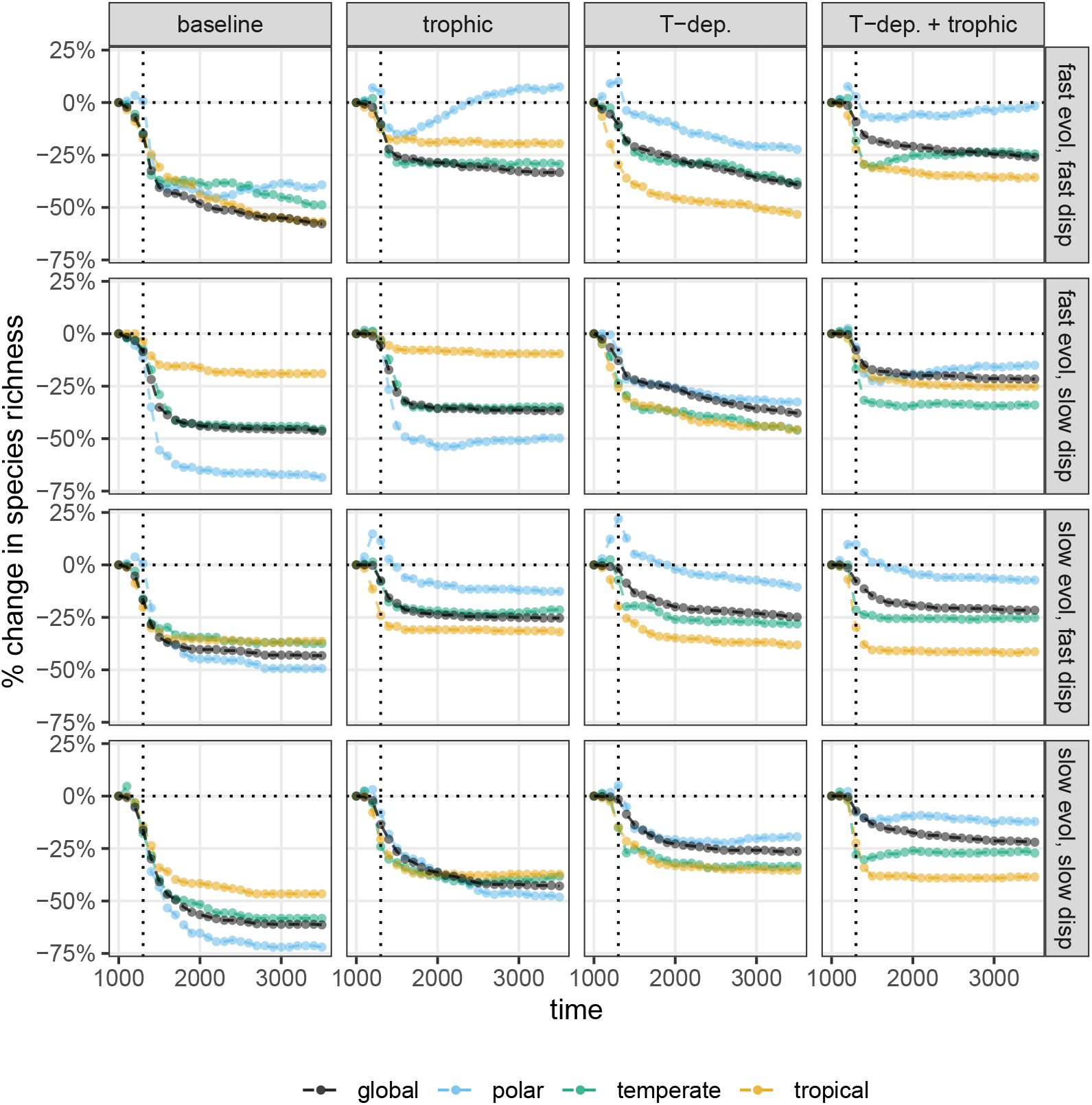
Relative change in global species richness from the community state at the onset of climate change (ordinate) over time (abscissa), averaged over replicates and given in 100-year steps (points). Black points correspond to species richness over the whole landscape; the blue points to richness in the top third of all patches (the polar region), green points to the middle third (temperate region), and yellow points to the last third (tropical region). Panel rows show different parameterizations (all four combinations of high and low genetic variance and dispersal ability); columns represent various model setups (the baseline model; an added trophic level of consumers; temperature-dependent competition coefficients; and the combined influence of consumers and temperature-dependent competition). Dotted horizontal lines highlight the point of no net change in global species richness; dotted vertical lines indicate the time at which climate change ends.

The identity of the species undergoing global extinction is not random, but strongly biased towards initially cold-adapted species. Indeed, looking at the effects of climate change on the fraction of patches occupied by species over the landscape reveals that initially cold-adapted species lose suitable habitat during climate change, and even afterwards (Figure 4). For the northernmost species, this always happens to the point where all habitat is lost, resulting in their extinction. While this pattern holds universally in every model setup and parameterization, the converse (range expansion) happens only under highly restrictive conditions, requiring both good dispersal ability and sufficient available genetic variance, as well as consumer pressure (Figure 4). Even so, only initially warm-adapted species expand their ranges; cold-adapted ones are still affected negatively, and as severely as in all other scenarios.

**Figure 4:**
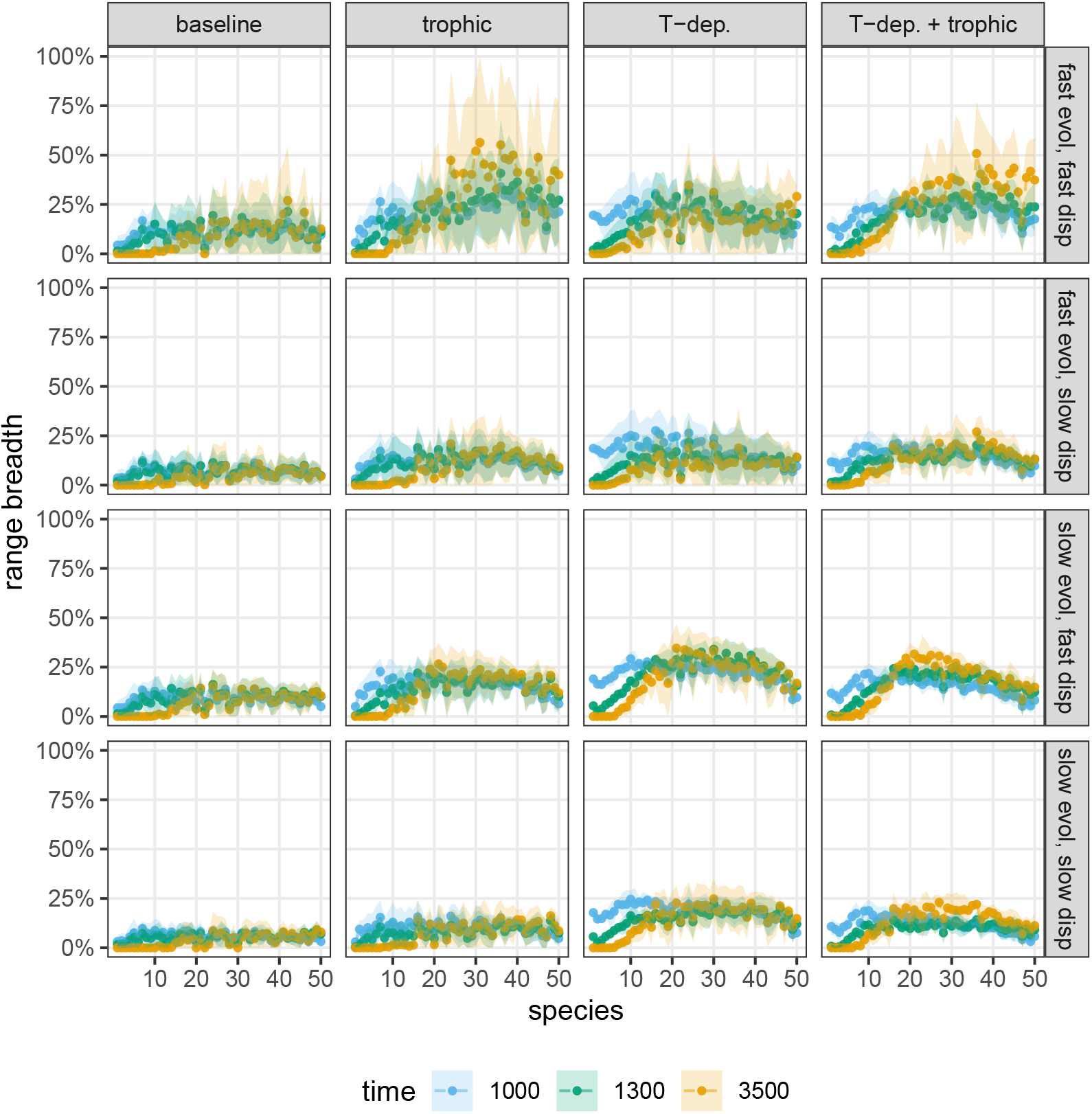
Range breadth of each species expressed as the percentage of the whole landscape they occupy (ordinate) at three different time stamps (colors). The mean (points) and plus/minus one standard deviation range (colored bands) are shown over replicates. Numbers along the abscissa represent species, with initially more warm-adapted species corresponding to higher values. The range breadth of each species is shown at three time stamps: at the start of climate change (1000 years, blue), the end of climate change (1300 years, green), and at the end of our simulations (3500 years, yellow). Panel layout as in Figure 3.

Comparing the baseline model with all the others in Figure 4, trophic interactions and temperature-dependent competition almost always increase species’ range breadths. Since this increase is also accompanied by fewer species disappearing overall (Figure 3), these mechanisms enhance local diversity by increasing the spatial overlap of species (Figure 5). The fostering of local coexistence by trophic interactions and temperature-dependent competition is in line with general ecological expectations. Predation pressure can enhance diversity by providing additional mechanisms of density regulation and thus prey coexistence through predator partitioning ^28,29^. In turn, temperature-dependent competition means species can reduce interspecific competition by evolving locally suboptimal mean temperature optima ^19^, compared with the baseline model’s fixed competition coefficients. An important question is how local diversity is affected when the two processes act simultaneously. In fact, any synergy between temperature-dependent competition and predation is very weak (Figure 5), and their combined contribution to local diversity patterns depends on the level of available genetic variance and ability to disperse.

**Figure 5:**
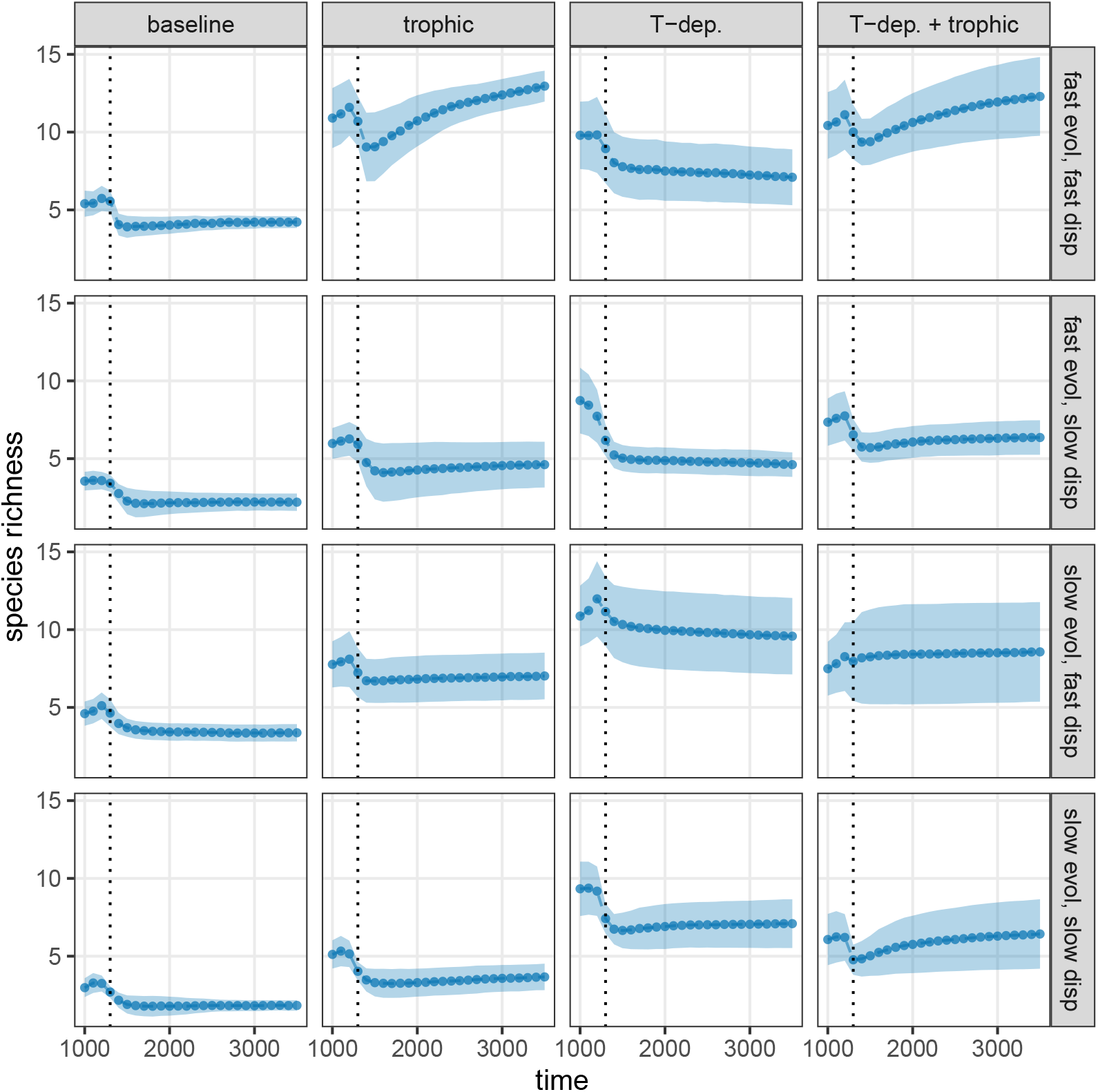
Local species richness of communities over time, from the start of climate change to the end of the simulation, averaged over replicates. Values are given in 100-year steps. At each point in time, the figure shows the mean number of species per patch over the landscape (points) and their standard deviation (shaded region, extending one standard deviation both up- and downwards from the mean). Panel layout as in Figure 3. Dotted vertical lines indicate the time at which climate change ends.

One can also look at larger regional changes in species richness, dividing the landscape into three equal parts: the top third (polar region), the middle third (temperate region), and the bottom third (tropical region). Region-wise exploration of changes in species richness (Figure 3) shows that the species richness of the polar region is highly volatile. It often experiences the greatest losses; however, with high dispersal ability, high genetic variance, and the presence of trophic interactions, regional richness can even increase compared to its starting level. Of course, change in regional species richness is a result of species dispersing to new patches and regions as well as of local extinctions. Since the initially most cold-adapted species lose their habitat and go extinct, altered regional species richness is connected to having altered community compositions along the spatial gradient. All regions experience turnover in species composition (SI, Section 5.1), but in general, the polar region experiences the largest turnover, where the final communities are at least 50% and sometimes more than 80% dissimilar to the community state right before the onset of climate change. This result is in agreement with previous studies ^7,30^; however, trophic interactions and temperature-dependent competition, both separately and in concert, reduce species turnover in all regions compared to the baseline model—a consequence of fewer species going globally extinct to begin with.

A further elucidating pattern is revealed by analyzing the relationship between the time-integrated temperature trait lag and community-wide trait dispersion (Figure 6). A strong negative correlation between the two shows the positive effect of species having more varied temperature tolerance strategies on the community’s ability to respond to climate change, as predicted by trait-driver theory ^25^. Communities generated by the four different model setups form more or less distinct clusters along this relationship. In comparison to the baseline model, trophic interactions cause a wider range of possible community-wide trait dispersion values, and thus a wider range of integrated trait lags. In contrast, temperature-dependent competition generally reduces community-wide trait dispersion, resulting in larger trait lags and thus less responsive communities. Interestingly, the same clustering of results is absent with respect to model parameterization in terms of available genetic variance and dispersal ability. This suggests that the relationship does not depend on these parameters, but is rather the property of the model as a whole. To check this, we have extended our simulations to a much wider range of parameter values, with average dispersal rates between 10^−3^ m/yr–10^4^ m/yr and average genetic variances between 10^−5 *◦*^C^2^–0.5 ^*◦*^C^2^ (from the original 10^−2^ m/yr–10^2^ m/yr and 10^−3 *◦*^C^2^–0.1 ^*◦*^C^2^; SI, Section 6). We find that the negative relationship is maintained, with the same slope and intercept as before (SI, Figure S4). Thus, changing these parameters does not affect the relationship between trait dispersion and trait lag.

**Figure 6:**
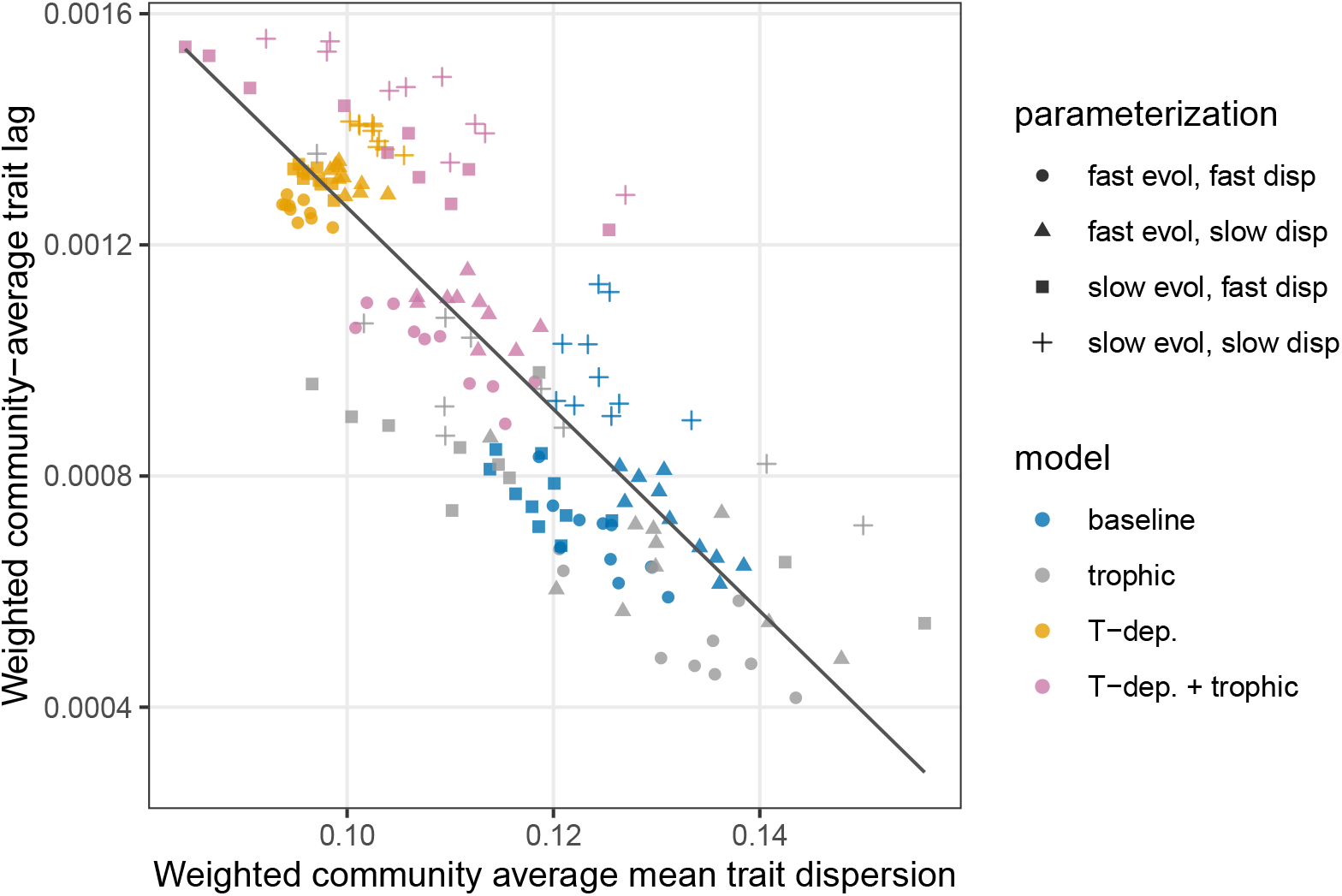
The ability of communities (points) to track local climatic conditions (ordinate), against observed trait diversity within those communities (abscissa). Both quantities are averaged over the landscape and time from the beginning to the end of the climate change period, yielding a single number for every community. The greater the average local diversity of mean temperature optima in a community, the closer it is able to match the prevailing temperature conditions (regression line; slope −0.017, intercept 0.003, *p <* 2.2 × 10^−16^). Species’ dispersal ability and available genetic variance (point symbols) are distributed evenly along the relationship, determining its slope and intercept. However, communities generated by different models (colors) are clustered: models with temperature-dependent competition have lower community-wide trait variation and thus a greater mismatch of their mean temperature optima with prevailing conditions than models without temperature-dependent competition.

## Discussion

This work introduced a modeling framework combining dispersal, evolutionary dynamics, and ecological interactions in a way that is tractable, easy to implement, fast to execute on a computer, and can handle ecological interactions of realistic complexity without simultaneously breaking other aspects of the approach. Individual-based models ^6^, for instance, do in principle allow one to include arbitrary levels of complexity, but tend to be computationally expensive. Other models yield detailed projections of individual species and their genetic structure but ignore species interactions altogether ^31^. An intermediate approach is based on quantitative genetics, which takes species interactions into account and yields a description of species’ genetic structure that is sufficiently simplified to be tractable. Earlier models in this spirit ^7,32^ were built on coupled partial differential equations. Unfortunately, while the theory behind such models is highly elegant, coupled nonlinear partial differential equations are notoriously difficult to implement in a way that is numerically stable, yields accurate results, and does not require unacceptably long run-times—notably, naive discretization schemes often do not work well. Unfortunately, despite persistent warnings about these problems ^33^, such naive solution schemes still prevail in the literature.

We circumvented this problem by building, from first principles, a different framework for spatial eco-evolutionary dynamics. Within a single patch, it is based on a quantitative genetic recursion model ^16,19^. Spatial locations are discretized from the outset, therefore the approach is built on ordinary differential equations alone. As a consequence, it executes fast even with substantial model complexity: on an ordinary desktop computer, a single run for 3500 years with both trophic interactions and temperature-dependent competition, with 50 species on both trophic levels and 50 habitat patches (for 50 × 100 × 2 = 10000 dynamical variables; the factor of 2 is because both the density and trait mean of each species may change) finished in less than 20 minutes. Incorporation of further model complexity is straightforward: complex food webs and spatial structure, further trait variables under selection (e.g., having both temperature optimum and body size evolve, the latter dictating the type of prey a species can consume ^34^) or an improved climate model with annual fluctuations and more detailed predictions of temperature profiles, can all be implemented. There is one important thing our model currently cannot do: since trait distributions are assumed to be normal with constant variance, a species cannot split into two daughter lineages in response to disruptive selection, as this would require the trait distribution to become gradually more and more bimodal. As such, our model ignores speciation, which may turn out to be an important process in regions which become species-impoverished following climate change.

Using this framework, we demonstrate that biotic factors such as trophic interactions and temperature-dependent competition are important in shaping species’ eco-evolutionary response to climate change—in fact, they can be as influential as the ability of species to adapt to new local climates or to disperse to new habitats. With trophic interactions and temperature-dependent competition, species have broader distributions, coexist to a higher degree, and global biodiversity loss is less severe, in comparison to the baseline model without the aforementioned dynamics. The importance of biotic interactions for shaping species’ response to climate change is well-known ^8,13,14^. Our work complements these studies by further demonstrating the significance of biotic interactions in an eco-evolutionary setting as well. The mechanisms behind this are predator-mediated coexistence ^28,29^ (in the case of trophic interactions), and reduced interspecific competition with increasing trait distance ^19^. Note that this last mechanism is not guaranteed to promote diversity, since the level of difference in mean temperature optima required for significant reductions in competition might mean that species have local growth rates that are too suboptimal for persistence. Thus, the ability of this mechanism to boost diversity depends on whether species are able to tolerate suboptimal climates sufficiently to avoid local competition.

Besides the importance of biotic interactions affecting species’ persistence and distribution under climate change, we also show that their dispersal ability and available genetic variance (i.e., capacity to respond to selective pressures swiftly) influence their responses. When local conditions change and temperatures increase, species become increasingly maladapted at their initial locations and pre-adapted to temperatures at higher latitudes, driving a northward movement. Dispersal is therefore suggested as a mechanism that provides spatial insurance to species ^35,36^, mitigating the negative impacts of climate change. However, a northward movement of initially warm-adapted species comes at the expense of the species located in the coldest regions which cannot disperse further ^30^, a consequence of dispersal that has been shown in previous studies ^7^. One might think that combining good dispersal ability with large genetic variance should temper this problem by allowing the northernmost species to adapt locally, and thus alleviate the negative impacts of increased temperatures better than each of these processes on their own. This expectation is also consistent with recent projections based on empirical data ^37^. However, the projected extinctions, considering both dispersal and species’ ability to adapt, have been obtained without explicitly considering species interactions ^37^. We show that large genetic variance combined with good dispersal ability result in a global biodiversity loss as high as when both dispersal ability and evolutionary rate are low. The reason, again, has to do with species interactions: the ability of individual species to disperse and adapt to new local conditions is of no use if they are prevented by other species from reaching the new locations. Similarly, cold-adapted species may be able to sustain in their current location with large genetic variance, but get out-competed by the arrival of better adapted migrating species. The surprising negative interaction between high dispersal and fast adaptation under climate change has also been demonstrated by Thompson and Fronhofer ^6^. However, in our case, we show that temperature-dependent competition reduces some of the negative impacts by allowing more local coexistence, albeit at the cost of reduced local growth rates.

Species’ traits are determined both by their evolutionary history and current selection pressures, constraining their possible values. Thus, the relationship between species richness and the distribution of species’ traits is not necessarily given. For example, one can have many species all with similar temperature optima (such as for corals) or one can have fewer species with a wider range of optima, like for phytoplankton in temperate lakes. The response to a specific environmental driver, such as temperature, depends more on the relevant community-wide trait dispersion than on species richness *per se*, but these two variables can potentially be statistically correlated, as often found in biodiversity and ecosystem functioning studies ^38^. Furthermore, other traits affecting community dynamics may be correlated with the trait that is affected by an environmental driver, such as temperature-dependent species interactions as demonstrated in this work. Despite this, it is encouraging for the general predictability of biotic climate impact models that the resulting trait dispersion in temperature-related traits strongly correlates with the ability of the community to cope with climate change. This can justify an argument that focus needs to be on processes that can sustain local community-wide trait dispersion, providing an argument for general biodiversity-enhancing measures such as preserving habitat heterogeneity, maintaining populations of keystone species, and for constructing dispersal corridors.

Biological communities are affected by many factors, ecological as well as evolutionary, which influence their response to climate change. Our framework demonstrates the importance of including relevant biological processes for predicting large-scale consequences of climate change on global and local biodiversity. Realistic mechanisms such as species interactions over multiple trophic levels and temperature-dependent competition, as well as particular combinations of dispersal and available genetic variance, can alleviate some of the negative impact of climate change, showing potential ways for ecological communities to adjust to altered climatic conditions. Despite this, the negative impact of climate change on ecological communities is severe, with numerous global extinctions and effects that are manifested long after the climate has again stabilized.

## Methods

We consider a chain of *L* evenly spaced patches along a latitudinal gradient, where patches 1 and *L* correspond to north pole and equator, respectively. The temperature *T^k^*(*t*) in patch *k* at time *t* is given by

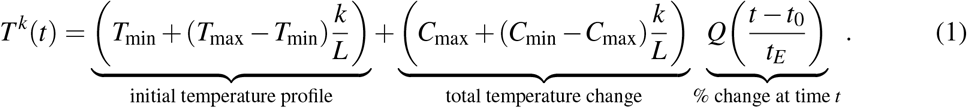

*T*_min_ and *T*_max_ are the initial polar and equatorial temperatures; *C*_max_ and *C*_min_ are the corresponding temperature increases after *t_E_* = 300 years, based on the IPCC intermediate emission scenario ^24^. *t*_0_ = 1000 years is an establishment time preceding climate change. *Q*(*τ*) describes the sigmoidal temperature increase in time: *Q*(*τ*) equals 0 for *τ <* 0, 1 for *τ >* 1, and 10*τ*^3^ 15*τ*^4^ + 6*τ*^5^ otherwise. Figure 2**B** depicts the resulting temperature change profile.

Combining quantitative genetics with dispersal across the *L* patches, we track the population density and mean temperature optimum of *S* species. Let 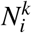 be the density and 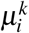 the mean temperature optimum of species *i* in patch *k* (subscripts denote species; superscripts patches). The governing equations then read

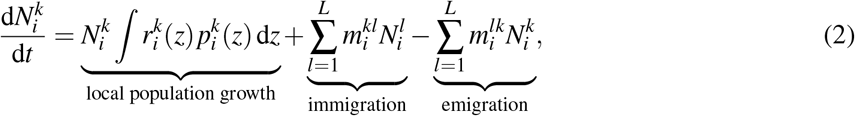

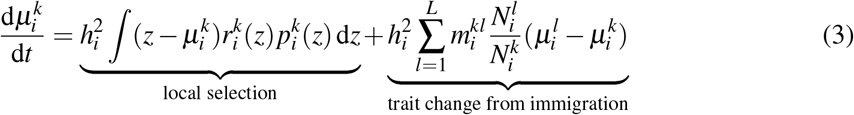

(SI, Section 1), where *t* is time, 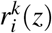 the per capita growth rate of species *i*’s phenotype *z* in patch *k*, 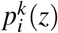 species *i*’s temperature optimum distribution in patch *k* (which is normal with patch-dependent mean 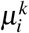 and patch-independent variance 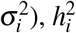 the heritability of species *i*’s temperature optimum, and 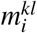 the migration rate of species *i* from patch *l* to *k*. The per capita growth rates 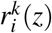 read

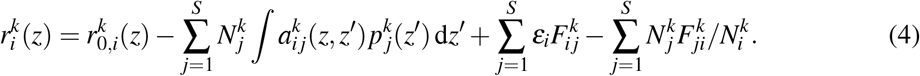

Here

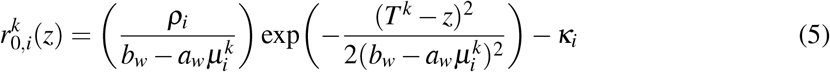

is the intrinsic growth of species *i*’s phenotype *z* in patch *k*. The constants *ρ_i_*, *b_w_*, and *a_w_* modulate a tradeoff between maximum growth and tolerance range ^23^ (Figure 2**A**), *κ_i_* is a mortality rate, and *T* ^*k*^ is the current local temperature in patch *k*. In turn, 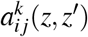 is the competition coefficient between species *i*’s phenotype *z* and species *j*’s phenotype *z*^/^ in patch *k*. We either assume constant, patch- and phenotype-independent coefficients *a_i j_*, or ones which decline with increasing trait differentiation according to

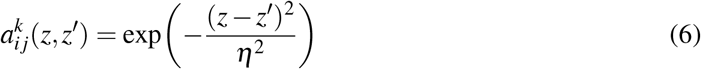

(temperature-dependent competition), where *η* is the competition width. The parameter *∊_i_* in Eq. 4 is species *i*’s resource conversion efficiency, and 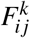 is the feeding rate of species *i* on *j* in patch *k*:

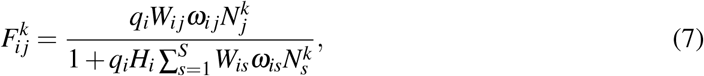

where *q_i_* is species *i*’s attack rate, *W_i j_* is the adjacency matrix of the feeding network (*W_i j_* = 1 if *i* eats *j* and 0 otherwise), *ω_i j_* is the proportion of effort of *i* on *j*, and *H_i_* is species *i*’s handling time. When adding a second trophic level, the number of species on the new level is equal to that at the lower level, and the feeding network is bipartite with connectance 0.5 (SI, Section 3.3).

We numerically integrated 10 replicates for each of 16 scenarios, made up of the fully factorial combinations of:

- The average dispersal rate between adjacent patches, which was either high (100 m/yr) or low (0.01 m/yr).
- The mean genetic variance per species, also either high (10^−1 *◦*^C^2^) or low (10^−3 *◦*^C^2^).
- The model setup, which was one of the following:
  1. One trophic level and constant competition coefficients, _*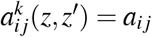*_.
  2. Two trophic levels and constant competition coefficients.
  3. One trophic level and competition coefficients given by Eq. 6.
  4. Two trophic levels and competition coefficients given by Eq. 6.

For each replicate, all other parameters are assigned based on Section 6 in the SI. Numerical integration of the system starts with initial conditions

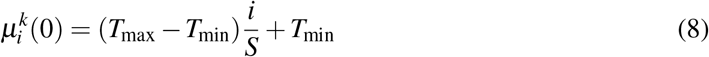

and

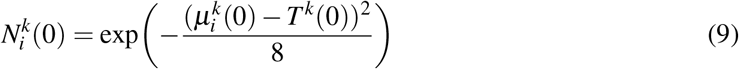

(SI, Section 3.7), and terminates after 3500 years.

The community-average trait dispersion 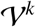 of the local community in patch *k* is the density-weighted variance of species’ mean temperature optima:

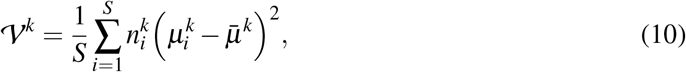

where 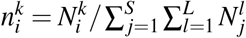 is the relative density of species *i* in patch *k*, and 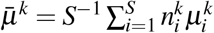 is the weighted average of species’ mean temperature optima in patch *k*. In turn, the community-average trait lag 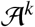 in patch *k* is defined as

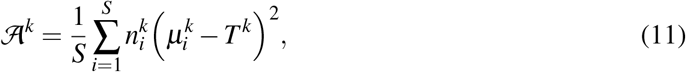

where *T*^*k*^ is the local temperature in patch *k*. In Figure 6, these quantities are averaged over all patches of the landscape and over time (from the beginning to the end of climate change). These averages are taken over each of the 160 model realizations (16 scenarios, with 10 replicates each).

## Acknowledgments

We would like to thank Priyanga Amarasekare and Peter Münger for discussions. This research was funded by the Swedish Research Council (grant FORMAS 2015-01262 to AE, and grant VR 2017-05245 to GB).

## Code availability

Computer code for implementing our model and replicating our results can be found at:
www.github.com/dysordys/spatial_ecoevo/

