## Supplementary Information for "The importance of species interactions in spatially explicit eco-evolutionary community dynamics under climate change"

#### 1 Quantitative genetic recursion in space

Let $S$ species be distributed across $L$ habitat patches, with possible migration in between. Each species $i$ is characterized by its density $N_i^k$ and trait distribution $p_i^k(z)$ in patch $k$. Here $z$ measures a unidimensional quantitative trait of interest; $N_i^k p_i^k(z)\mathrm{d}z$ is then the density of species $i$'s individuals in patch $k$ whose phenotype values fall between $z$ and $z + \mathrm{d}z$. By definition,

$$\int p_i^k(z)\,\mathrm{d}z = 1 \tag{S1}$$

at any moment of time. The mean trait of species $i$ in patch $k$ is

$$\mu_i^k = \int z p_i^k(z)\,\mathrm{d}z. \tag{S2}$$

Our starting point is the framework of quantitative genetic recursion (Lande 1976, Slatkin 1980, Taper and Case 1985, 1992, Barabás and D'Andrea 2016) in the weak selection limit (Bürger 2011). We extend this approach by adding migration. The basic equation giving the change in the density distribution of species $i$ in patch $k$ reads

$$\tilde{N}_i^k \tilde{p}_i^k(z) = W_i^k(z) N_i^k p_i^k(z) + \sum_{l=1}^{L} M_i^{kl} N_i^l p_i^l(z) - \sum_{l=1}^{L} M_i^{lk} N_i^k p_i^k(z), \tag{S3}$$

where the tilde denotes values after selection and migration, but before reproduction. The first term on the right hand side describes selection via the fitness function $W_i^k(z)$. The second term is immigration from all other patches (therefore $M_i^{kl}$ is the dispersal rate from patch $l$ to patch $k$, with $M_i^{kk} = 0$ for all $i$ and $k$). The final term is the loss of species $i$'s trait $z$ in patch $k$ due to emigration from the focal patch.

For the underlying genetics, we assume a large number of loci each contributing a very small additive effect to the trait (Barton et al. 2017, Turelli 2017). In this infinitesimal model, all trait distributions $p_i^k(z)$ are normal with a variance $\sigma_i^2$ that does not change in response to selection:

$$p_i^k(z) = \frac{1}{\sigma_i\sqrt{2\pi}} \exp\left(-\frac{(z-\mu_i^k)^2}{2\sigma_i^2}\right). \tag{S4}$$

The change in mean trait from one generation to the next is given by the breeder's equation (e.g., Falconer 1981):

$$\hat{\mu}_i^k - \mu_i^k = h_i^2(\tilde{\mu}_i^k - \mu_i^k), \tag{S5}$$

where the hat denotes values in the next generation, and $\tilde{\mu}_i^k$ denotes the trait mean after selection and migration, but before reproduction. After reproduction, $\hat{N}_i^k = \tilde{N}_i^k$, and $\hat{p}_i^k$ is given by Eq. S4 with $\hat{\mu}_i^k$ calculated from the breeder's equation (Barabás and D'Andrea 2016).

In the weak selection limit, the fitness $W_i^k(z)$ of species $i$'s phenotype $z$ in patch $k$ can be written

$$W_i^k(z) = 1 + s r_i^k(z), \tag{S6}$$

where $s$ is a small parameter, and $r_i^k(z)$ is the per capita growth rate of species $i$'s phenotype $z$ in patch $k$, defined by ecological interactions. The migration rates are similarly written as

$$M_i^{kl} = s m_i^{kl}. \tag{S7}$$

To obtain the dynamics of the densities, we integrate Eq. S3 across $z$:

$$\tilde{N}_i^k \int \tilde{p}_i^k \,\mathrm{d}z = N_i^k \int W_i^k(z) p_i^k(z) \,\mathrm{d}z + \sum_{l=1}^{L} M_i^{kl} N_i^l \int p_i^l(z) \,\mathrm{d}z - \sum_{l=1}^{L} M_i^{lk} N_i^k \int p_i^k(z) \,\mathrm{d}z. \tag{S8}$$

Using Eq. S1 and $\tilde{N}_i^k = \hat{N}_i^k$:

$$\hat{N}_i^k = N_i^k \int W_i^k(z) p_i^k(z) \,\mathrm{d}z + \sum_{l=1}^{L} M_i^{kl} N_i^l - \sum_{l=1}^{L} M_i^{lk} N_i^k. \tag{S9}$$

In the weak selection limit (Eqs. S6 and S7),

$$\hat{N}_i^k = N_i^k \int \left[1 + s r_i^k(z)\right] p_i^k(z) \,\mathrm{d}z + s \sum_{l=1}^{L} m_i^{kl} N_i^l - s \sum_{l=1}^{L} m_i^{lk} N_i^k, \tag{S10}$$

which, using Eq. S1, is written as

$$\hat{N}_i^k = N_i^k + s N_i^k \int r_i^k(z) p_i^k(z) \,\mathrm{d}z + s \sum_{l=1}^{L} m_i^{kl} N_i^l - s \sum_{l=1}^{L} m_i^{lk} N_i^k. \tag{S11}$$

Subtracting $N_i^k$ and dividing by $s$ leads to

$$\frac{\hat{N}_i^k - N_i^k}{s} = N_i^k \int r_i^k(z) p_i^k(z) \,\mathrm{d}z + \sum_{l=1}^{L} m_i^{kl} N_i^l - \sum_{l=1}^{L} m_i^{lk} N_i^k. \tag{S12}$$

With appropriate scaling (Barabás and D'Andrea 2016), this can be written in differential equation form when $s$ is sufficiently small:

$$\frac{\mathrm{d}N_i^k}{\mathrm{d}t} = N_i^k \int r_i^k(z) p_i^k(z) \,\mathrm{d}z + \sum_{l=1}^{L} m_i^{kl} N_i^l - \sum_{l=1}^{L} m_i^{lk} N_i^k. \tag{S13}$$

To obtain the equation for the change in trait means, we write $\tilde{\mu}_i^k$ by rearranging Eq. S3:

$$\tilde{p}_i^k = \frac{W_i^k(z) N_i^k p_i^k(z) + \sum_{l=1}^{L} M_i^{kl} N_i^l p_i^l(z) - \sum_{l=1}^{L} M_i^{lk} N_i^k p_i^k(z)}{\hat{N}_i^k}. \tag{S14}$$

We use Eq. S9 to expand the denominator:

$$\tilde{p}_i^k = \frac{W_i^k(z) N_i^k p_i^k(z) + \sum_{l=1}^{L} M_i^{kl} N_i^l p_i^l(z) - \sum_{l=1}^{L} M_i^{lk} N_i^k p_i^k(z)}{N_i^k \int W_i^k(z) p_i^k(z) \,\mathrm{d}z + \sum_{l=1}^{L} M_i^{kl} N_i^l - \sum_{l=1}^{L} M_i^{lk} N_i^k}. \tag{S15}$$

We multiply both sides by $z$ and integrate. Using Eq. S2, we get

$$\tilde{\mu}_i^k = \frac{N_i^k \int z W_i^k(z) p_i^k(z)\,\mathrm{d}z + \sum_{l=1}^{L} M_i^{kl} N_i^l \mu_i^l - \sum_{l=1}^{L} M_i^{lk} N_i^k \mu_i^k}{N_i^k \int W_i^k(z) p_i^k(z)\,\mathrm{d}z + \sum_{l=1}^{L} M_i^{kl} N_i^l - \sum_{l=1}^{L} M_i^{lk} N_i^k}. \tag{S16}$$

In the weak selection limit (Eqs. S6 and S7), this reads

$$\tilde{\mu}_i^k = \frac{N_i^k \int z(1+s r_i^k(z)) p_i^k(z)\,\mathrm{d}z + s\sum_{l=1}^{L} m_i^{kl} N_i^l \mu_i^l - s\sum_{l=1}^{L} m_i^{lk} N_i^k \mu_i^k}{N_i^k \int (1+s r_i^k(z)) p_i^k(z)\,\mathrm{d}z + s\sum_{l=1}^{L} m_i^{kl} N_i^l - s\sum_{l=1}^{L} m_i^{lk} N_i^k}. \tag{S17}$$

Using Eqs. S1 and S2, we get

$$\tilde{\mu}_i^k = \frac{N_i^k \mu_i^k + s N_i^k \int z r_i^k(z) p_i^k(z)\,\mathrm{d}z + s\sum_{l=1}^{L} m_i^{kl} N_i^l \mu_i^l - s\sum_{l=1}^{L} m_i^{lk} N_i^k \mu_i^k}{N_i^k + s N_i^k \int r_i^k(z) p_i^k(z)\,\mathrm{d}z + s\sum_{l=1}^{L} m_i^{kl} N_i^l - s\sum_{l=1}^{L} m_i^{lk} N_i^k}. \tag{S18}$$

Simplifying by $N_i^k$ and factoring out $s$ from both the numerator and denominator:

$$\tilde{\mu}_i^k = \frac{\mu_i^k + s\left[\int z r_i^k(z) p_i^k(z)\,\mathrm{d}z + \sum_{l=1}^{L} m_i^{kl} \frac{N_i^l}{N_i^k} \mu_i^l - \sum_{l=1}^{L} m_i^{lk} \mu_i^k\right]}{1 + s\left[\int r_i^k(z) p_i^k(z)\,\mathrm{d}z + \sum_{l=1}^{L} m_i^{kl} \frac{N_i^l}{N_i^k} - \sum_{l=1}^{L} m_i^{lk}\right]}. \tag{S19}$$

Taylor expanding around $s = 0$ to first order, we get

$$\begin{aligned} \tilde{\mu}_i^k = \mu_i^k + s\Bigg[&\int z r_i^k(z) p_i^k(z)\,\mathrm{d}z + \sum_{l=1}^{L} m_i^{kl} \frac{N_i^l}{N_i^k} \mu_i^l - \sum_{l=1}^{L} m_i^{lk} \mu_i^k \\ &-\mu_i^k\left(\int r_i^k(z) p_i^k(z)\,\mathrm{d}z + \sum_{l=1}^{L} m_i^{kl} \frac{N_i^l}{N_i^k} - \sum_{l=1}^{L} m_i^{lk}\right)\Bigg] + \mathscr{O}(s^2), \end{aligned} \tag{S20}$$

where $\mathscr{O}(s^2)$ means terms of order $s^2$ or higher. Simplification yields

$$\tilde{\mu}_i^k = \mu_i^k + s\left[\int (z-\mu_i^k) r_i^k(z) p_i^k(z)\,\mathrm{d}z + \sum_{l=1}^{L} m_i^{kl} \frac{N_i^l}{N_i^k} (\mu_i^l - \mu_i^k)\right] + \mathscr{O}(s^2). \tag{S21}$$

Substituting this expression into the breeder's equation (Eq. S5):

$$\hat{\mu}_i^k - \mu_i^k = h_i^2 s\left[\int (z-\mu_i^k) r_i^k(z) p_i^k(z)\,\mathrm{d}z + \sum_{l=1}^{L} m_i^{kl} \frac{N_i^l}{N_i^k} (\mu_i^l - \mu_i^k)\right]. \tag{S22}$$

After dividing both sides by $s$ and performing the $s \to 0$ limit with appropriate scaling (Barabás and D'Andrea 2016), we get the differential equation approximation

$$\frac{\mathrm{d}\mu_i^k}{\mathrm{d}t} = h_i^2\left[\int (z-\mu_i^k) r_i^k(z) p_i^k(z)\,\mathrm{d}z + \sum_{l=1}^{L} m_i^{kl} \frac{N_i^l}{N_i^k} (\mu_i^l - \mu_i^k)\right]. \tag{S23}$$

#### 2 Per capita growth rates and species-level dynamics

We model a trophic community in space, with $L$ distinct habitat patches. Each patch experiences an average temperature, which is increasing through time due to climate change. The evolvable trait $z$ is identified with temperature tolerance: an individual with trait $z$ achieves maximal intrinsic growth in a habitat with temperature $z$. Local population dynamics, excluding migration, are governed by four processes: temperature-dependent intrinsic growth, intra- and interspecific competition, growth due to consumption, and loss due to being consumed. The per capita growth rate of of species $i$'s phenotype $z$ in patch $k$ is written

$$r_i^k(z) = r_{0,i}^k(z) - \sum_{j=1}^{S} N_j^k \int a_{ij}^k(z,z') p_j^k(z') \,\mathrm{d}z' + \sum_{j=1}^{S} \varepsilon_i F_{ij}^k - \sum_{j=1}^{S} N_j^k F_{ji}^k / N_i^k, \tag{S24}$$

where, in patch $k$, $r_{0,i}^k(z)$ is the intrinsic growth rate of species $i$'s phenotype $z$, $a_{ij}^k(z,z')$ are competition coefficients between species $i$'s phenotype $z$ and species $j$'s phenotype $z'$, $\varepsilon_i$ is species $i$'s resource conversion efficiency, and $F_{ij}^k$ is the feeding rate of species $i$ on $j$. It has the form

$$F_{ij}^k = \frac{q_i W_{ij} \omega_{ij} N_j^k}{1 + q_i H_i \sum_{s=1}^{S} W_{is} \omega_{is} N_s^k}, \tag{S25}$$

where $q_i$ is species $i$'s attack rate, $W_{ij}$ is the adjacency matrix of the feeding network ($W_{ij} = 1$ if $i$ eats $j$ and 0 otherwise), $\omega_{ij}$ is the relative consumption rate of $i$ on $j$ (we assume an equal split across resources, so that $\omega_{ij} = [\text{number of resources}]^{-1}$ for all consumers), and $H_i$ is species $i$'s handling time.

Substituting the growth rate in Eq. S24 into Eqs. S13 and S23 yields

$$\begin{aligned} \frac{\mathrm{d}N_i^k}{\mathrm{d}t} &= N_i^k \int r_{0,i}^k(z) p_i^k(z) \,\mathrm{d}z - N_i^k \int \sum_{j=1}^{S} N_j^k \left[ \int a_{ij}^k(z,z') p_j^k(z') \,\mathrm{d}z' \right] p_i^k(z) \,\mathrm{d}z \\ &\quad + N_i^k \int \left[ \sum_{j=1}^{S} \varepsilon_i F_{ij}^k - \sum_{j=1}^{S} N_j^k F_{ji}^k / N_i^k \right] p_i^k(z) \,\mathrm{d}z + \sum_{l=1}^{L} m_i^{kl} N_i^l - \sum_{l=1}^{L} m_i^{lk} N_i^k \end{aligned} \tag{S26}$$

for the population densities, and

$$\begin{aligned} \frac{\mathrm{d}\mu_i^k}{\mathrm{d}t} &= h_i^2 \int (z - \mu_i^k) r_{0,i}^k(z) p_i^k(z) \,\mathrm{d}z - h_i^2 \int (z - \mu_i^k) \sum_{j=1}^{S} N_j^k \left[ \int a_{ij}^k(z,z') p_j^k(z') \,\mathrm{d}z' \right] p_i^k(z) \,\mathrm{d}z \\ &\quad + h_i^2 \int \left[ \sum_{j=1}^{S} \varepsilon_i F_{ij}^k - \sum_{j=1}^{S} N_j^k F_{ji}^k / N_i^k \right] (z - \mu_i^k) p_i^k(z) \,\mathrm{d}z + h_i^2 \sum_{l=1}^{L} m_i^{kl} \frac{N_i^l}{N_i^k} (\mu_i^l - \mu_i^k) \end{aligned} \tag{S27}$$

for the trait means. Rearranging, we get

$$\begin{aligned} \frac{\mathrm{d}N_i^k}{\mathrm{d}t} &= N_i^k \int r_{0,i}^k(z) p_i^k(z) \,\mathrm{d}z - N_i^k \sum_{j=1}^{S} N_j^k \iint p_i^k(z) a_{ij}^k(z,z') p_j^k(z') \,\mathrm{d}z' \mathrm{d}z \\ &\quad + N_i^k \left[ \sum_{j=1}^{S} \varepsilon_i F_{ij}^k - \sum_{j=1}^{S} N_j^k F_{ji}^k / N_i^k \right] \int p_i^k(z) \,\mathrm{d}z + \sum_{l=1}^{L} m_i^{kl} N_i^l - \sum_{l=1}^{L} m_i^{lk} N_i^k, \end{aligned} \tag{S28}$$

$$\begin{aligned} \frac{\mathrm{d}\mu_i^k}{\mathrm{d}t} &= h_i^2 \int (z - \mu_i^k) r_{0,i}^k(z) p_i^k(z) \,\mathrm{d}z - h_i^2 \sum_{j=1}^{S} N_j^k \iint (z - \mu_i^k) p_i^k(z) a_{ij}^k(z,z') p_j^k(z') \,\mathrm{d}z' \mathrm{d}z \\ &\quad + h_i^2 \left[ \sum_{j=1}^{S} \varepsilon_i F_{ij}^k - \sum_{j=1}^{S} N_j^k F_{ji}^k / N_i^k \right] \int (z - \mu_i^k) p_i^k(z) \,\mathrm{d}z + h_i^2 \sum_{l=1}^{L} m_i^{kl} \frac{N_i^l}{N_i^k} (\mu_i^l - \mu_i^k). \end{aligned} \tag{S29}$$

By Eq. S1, the integral in the third term of Eq. S28 is simply 1. In turn, the integral in the third term of Eq. S29 is zero, because the integrand is a product of an odd and an even function in $z-\mu_i^k$. This leads to

$$\frac{\mathrm{d}N_i^k}{\mathrm{d}t} = N_i^k \int r_{0,i}^k(z) p_i^k(z)\,\mathrm{d}z - N_i^k \sum_{j=1}^{S} N_j^k \iint p_i^k(z) a_{ij}^k(z,z') p_j^k(z')\,\mathrm{d}z'\mathrm{d}z \tag{S30}$$

$$+ \sum_{j=1}^{S} \varepsilon_i N_i^k F_{ij}^k - \sum_{j=1}^{S} N_j^k F_{ji}^k + \sum_{l=1}^{L} m_i^{kl} N_i^l - \sum_{l=1}^{L} m_i^{lk} N_i^k,$$

$$\frac{\mathrm{d}\mu_i^k}{\mathrm{d}t} = h_i^2 \int (z-\mu_i^k) r_{0,i}^k(z) p_i^k(z)\,\mathrm{d}z - h_i^2 \sum_{j=1}^{S} N_j^k \iint (z-\mu_i^k) p_i^k(z) a_{ij}^k(z,z') p_j^k(z')\,\mathrm{d}z'\mathrm{d}z \tag{S31}$$

$$+ h_i^2 \sum_{l=1}^{L} m_i^{kl} \frac{N_i^l}{N_i^k} (\mu_i^l - \mu_i^k).$$

Introducing the definitions

$$b_i^k = \int r_{0,i}^k(z) p_i^k(z)\,\mathrm{d}z, \tag{S32}$$

$$\alpha_{ij}^k = \iint p_i^k(z) a_{ij}^k(z,z') p_j^k(z')\,\mathrm{d}z'\mathrm{d}z, \tag{S33}$$

$$g_i^k = \int (z-\mu_i^k) r_{0,i}^k(z) p_i^k(z)\,\mathrm{d}z, \tag{S34}$$

$$\beta_{ij}^k = \iint (z-\mu_i^k) p_i^k(z) a_{ij}^k(z,z') p_j^k(z')\,\mathrm{d}z'\mathrm{d}z, \tag{S35}$$

Eqs. S30-S31 read

$$\frac{\mathrm{d}N_i^k}{\mathrm{d}t} = N_i^k b_i^k - N_i^k \sum_{j=1}^{S} \alpha_{ij}^k N_j^k + \sum_{j=1}^{S} \varepsilon_i N_i^k F_{ij}^k - \sum_{j=1}^{S} N_j^k F_{ji}^k + \sum_{l=1}^{L} m_i^{kl} N_i^l - N_i^k \sum_{l=1}^{L} m_i^{lk}, \tag{S36}$$

$$\frac{\mathrm{d}\mu_i^k}{\mathrm{d}t} = h_i^2 g_i^k - h_i^2 \sum_{j=1}^{S} \beta_{ij}^k N_j^k + h_i^2 \sum_{l=1}^{L} m_i^{kl} \frac{N_i^l}{N_i^k} (\mu_i^l - \mu_i^k), \tag{S37}$$

with $F_{ij}^k$ given by Eq. S25.

### 3 Model parameterization

Here we present the detailed derivation and parameterization of each model component. Numerical values of the model's parameters, along with their units and short descriptions, are collected in Tables S1-S2.

#### 3.1 Temperature-dependent intrinsic growth rates

Following Amarasekare and Johnson (2017), we model the temperature-dependence of $r_{0,i}^k(z)$ using the Gaussian form

$$r_{0,i}^k(z) = A_i^k \exp\left(-\frac{(T^k - z)^2}{2(w_i^k)^2}\right) - \kappa_i, \tag{S38}$$

where $\kappa_i$ is a mortality rate for species $i$, $A_i^k - \kappa_i$ is the maximum intrinsic growth that can be achieved for species $i$ in patch $k$, $T^k$ is the temperature in patch $k$, and $w_i^k$ is the species- and patch-specific width of the growth curve.

The $w_i^k$ are not constant, but coevolve with species' mean temperature optima $\mu_i^k$. Within reasonable limits (such that the tolerance widths remain positive), this can be modeled using the following linear relationship:

$$w_i^k = b_w - a_w \mu_i^k, \tag{S39}$$

where $a_w$ and $b_w$ are positive constant parameters. This creates a trend of increasingly broader curves for lower temperature optima, corresponding to the observed correlation between tolerance width and latitude (Addo-Bediako et al. 2000, Sunday et al. 2011). For determining $A_i^k$, we use the observation that there is a tradeoff between the width and maximum height of temperature optima. This phenomenon is brought about by higher annual temperature variability at higher latitudes: since reaction norms are nonlinear functions of temperature, the optimal response to the mean temperature is not the same as the mean optimal response, due to Jensen's inequality (Amarasekare and Johnson 2017). This also means that the distance between experienced and optimal temperatures is larger at higher latitudes, where climate tends to be more variable (Deutsch et al. 2008). However, our model does not take annual temperature fluctuations into account, which means that the observed optimum shift cannot evolve as in Amarasekare and Johnson (2017). Instead, we imposed this tradeoff by assigning a smaller $A_i^k$ to wider temperature adaptations at low $\mu_i^k$ values (Eq. S39), mimicking the fact that populations at higher latitudes are adapted to more variable environments and thus also living at temperatures further away from those that would yield maximum growth in a stable environment. The tradeoff is implemented as

$$A_i^k = \frac{\rho_i}{w_i^k} = \frac{\rho_i}{b_w - a_w \mu_i^k}, \tag{S40}$$

where $\rho_i$ is a parameter modulating the shape of the tradeoff. Substituting the parameter specifications of Eqs. S39-S40 into Eq. S38 gives

$$r_{0,i}^k(z) = \left( \frac{\rho_i}{b_w - a_w \mu_i^k} \right) \exp\left( -\frac{(T^k - z)^2}{2(b_w - a_w \mu_i^k)^2} \right) - \kappa_i. \tag{S41}$$

This form of the tradeoff, and its particular parameterization (Table S1), were chosen so the resulting growth curves would qualitatively mimick those found empirically (e.g., Deutsch et al. 2008). Figure S1 shows what the growth rates of Eq. S41 look like, in six species equally spaced along a temperature gradient, both for consumers and resources.

To obtain the species-level parameters $b_i^k$ and $g_i^k$ from this growth rate, we substitute Eqs. S4 and S41 into Eqs. S32 and S34:

$$b_i^k = \int \left[ \left( \frac{\rho_i}{b_w - a_w \mu_i^k} \right) \exp\left( -\frac{(T^k - z)^2}{2(b_w - a_w \mu_i^k)^2} \right) - \kappa_i \right] \frac{\exp\left( -\frac{(z-\mu_i^k)^2}{2\sigma_i^2} \right)}{\sigma_i \sqrt{2\pi}} \, \mathrm{d}z, \tag{S42}$$

$$g_i^k = \int (z - \mu_i^k) \left[ \left( \frac{\rho_i}{b_w - a_w \mu_i^k} \right) \exp\left( -\frac{(T^k - z)^2}{2(b_w - a_w \mu_i^k)^2} \right) - \kappa_i \right] \frac{\exp\left( -\frac{(z-\mu_i^k)^2}{2\sigma_i^2} \right)}{\sigma_i \sqrt{2\pi}} \, \mathrm{d}z. \tag{S43}$$

The integrals can be evaluated directly, leading to the final forms

$$b_i^k = \left( \frac{\rho_i}{b_w - a_w \mu_i^k} \right) \frac{b_w - a_w \mu_i^k}{\sqrt{(b_w - a_w \mu_i^k)^2 + \sigma_i^2}} \exp\left( -\frac{(T^k - \mu_i^k)^2}{2[(b_w - a_w \mu_i^k)^2 + \sigma_i^2]} \right) - \kappa_i, \tag{S44}$$

$$g_i^k = \left( \frac{\rho_i}{b_w - a_w \mu_i^k} \right) \frac{\sigma_i^2 (b_w - a_w \mu_i^k)(T^k - \mu_i^k)}{[(b_w - a_w \mu_i^k)^2 + \sigma_i^2]^{3/2}} \exp\left( -\frac{(T^k - \mu_i^k)^2}{2[(b_w - a_w \mu_i^k)^2 + \sigma_i^2]} \right). \tag{S45}$$

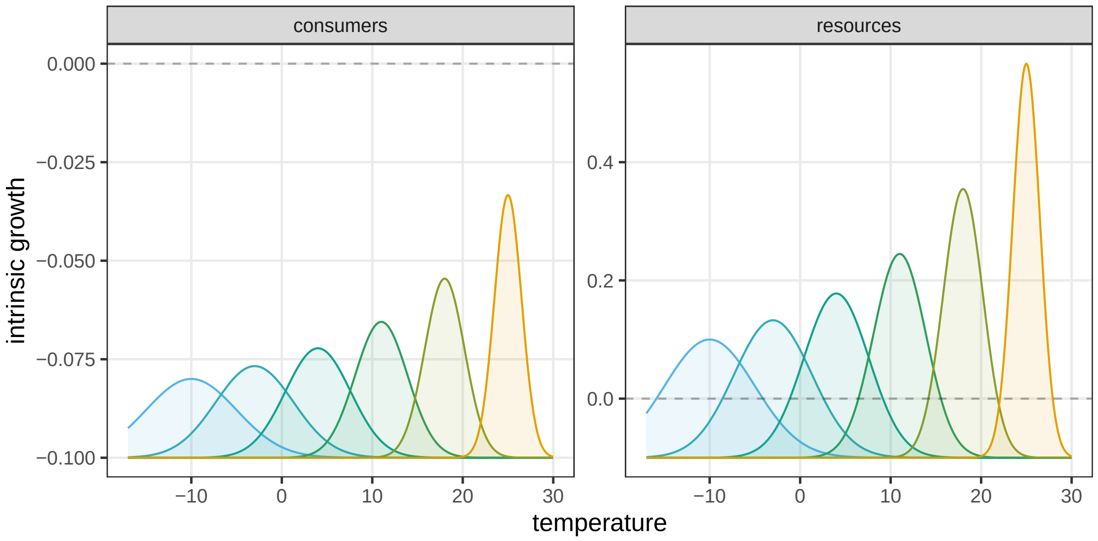


**Figure S1:** Tradeoff between the height and breadth of temperature optima, for phenotypes of six consumer (left) and six resource species (right). Colors indicate species, with cooler shades standing for more cold-adapted ones. The dashed line indicates zero growth; therefore the $r_{0,i}^k$ of consumer species is always a mortality rate, which must be compensated by consumption. For resources, $r_{0,i}^k$ is positive unless the local temperature is too far from the optimum for that species. In this figure, $\kappa_i = 0.1$ for all species, $\rho_i = 0.1$ for all consumers, and $\rho_i = 0.1$ for all resources. In our simulations, to make sure that the model's predicted community patterns are not an artifact of the above precise tradeoff shapes, the $\rho_i$ were randomized in a $\pm 10\%$ range of these values (Table S1).

#### 3.2 Competition coefficients

We only model competition between resource species. Consumers also compete but only indirectly, via shared resources. This means that the competition kernel $a_{ij}^k(z,z')$ is zero unless both $i$ and $j$ refer to resource species. When they do, we use two different kernel forms. One only depends on species identity, and is independent of trait or patch value:

$$a_{ij}^k(z,z') = a_{ij}. \tag{S46}$$

Applying Eqs. S33 and S35 to obtain $\alpha_{ij}$ and $\beta_{ij}$, we get

$$\alpha_{ij}^k = a_{ij} \iint p_i^k(z) p_j^k(z') \,\mathrm{d}z' \mathrm{d}z = a_{ij}, \tag{S47}$$

$$\beta_{ij}^k = a_{ij} \iint (z-\mu_i^k) p_i^k(z) p_j^k(z') \,\mathrm{d}z' \mathrm{d}z = 0. \tag{S48}$$

In the first equation, the integrals evaluate to 1 because of Eq. S4, while in the second equation, integration with respect to $z$ involves the product of an odd and an even function in $z - \mu_i^k$ and is therefore zero. The constant $a_{ij}$ values are sampled randomly (Table S1), in a way that ensures intraspecific competition being always stronger than interspecific competition.

The second form of the competition kernel we use depends on the difference between phenotypic values:

$$a_{ij}^k(z,z') = \exp\left(-\frac{(z-z')^2}{\eta^2}\right), \tag{S49}$$

where $\eta$ is the competition width, determining how distant two phenotypes must be for competition to be significantly reduced between them. Here $\alpha_{ij}^k$ and $\beta_{ij}^k$ can be calculated by direct integration using Eqs. S33 and S35:

$$\alpha_{ij}^k = \frac{\eta}{\sqrt{2\sigma_i^2 + 2\sigma_j^2 + \eta^2}} \exp\left( -\frac{(\mu_i^k - \mu_j^k)^2}{2\sigma_i^2 + 2\sigma_j^2 + \eta^2} \right), \tag{S50}$$

$$\beta_{ij}^k = -\frac{2\eta\sigma_i^2(\mu_i^k - \mu_j^k)}{(2\sigma_i^2 + 2\sigma_j^2 + \eta^2)^{3/2}} \exp\left( -\frac{(\mu_i^k - \mu_j^k)^2}{2\sigma_i^2 + 2\sigma_j^2 + \eta^2} \right). \tag{S51}$$

This form of the competition kernel facilitates trait divergence even within a single patch, since it may be profitable for a species to specialize on a suboptimal temperature to thus weaken interspecific competition and coexist locally. Biologically, this mechanism is explained by having microhabitats within each patch, with higher and lower local temperatures than the patch average. A species with a higher or lower temperature optimum than the average can exploit these microhabitats better than the locally dominant species, and therefore persist in the patch as a whole—though it will not be able to achieve as much growth due to the relatively limited availability of such microhabitats.

##### 3.3 Feeding network

We model two strict trophic levels, with $S$ resource and $S$ consumer species. The bipartite network of feeding connections $W_{ij}$ (with $W_{ij} = 1$ if $i$ eats $j$ and 0 otherwise) is generated as follows. First, both resources and consumers are labeled consecutively, based on their initial temperature adaptations: resource 1 / consumer 1 are the most cold-adapted, and resource $S$ / consumer $S$ the most warm-adapted. Next, we always put a feeding link between consumer $i$ and resource $i$. Finally, each consumer is randomly linked to $S/2 - 1$ other resources (rounded if necessary). For $S$ even, this will lead to the feeding network having a connectance of $1/2$.

##### 3.4 Genetic and environmental variances

Genetic variances $V_{G,i}$ are randomly drawn for each species, while the environmental variances $V_{E,i}$ are fixed (Table S2). We assume that there is no epistatic or dominance variance, which means that the total phenotypic variance, $\sigma_i^2$, is the sum of the additive genetic and environmental variances: $\sigma_i^2 = V_{G,i} + V_{E,i}$. The heritability of species $i$'s trait value is, by definition, the ratio of genetic to total phenotypic variance (e.g., Falconer 1981), so we have

$$h_i^2 = \frac{V_{G,i}}{V_{G,i} + V_{E,i}} = \frac{V_{G,i}}{\sigma_i^2}. \tag{S52}$$

Our model has no demographic stochasticity. This means we are assuming that even "low" population densities are sufficiently high to neglect genetic and ecological drift. This, however, can introduce problems with artifactual evolutionary rescue, where an unrealistically tiny population evolves to the point where it can finally grow, thus surviving and establishing itself even though it was bound to go extinct. To control for this artifact, we introduced a reduction in species' genetic variances whenever their local population densities dropped below the threshold $N_c$. Below this threshold, genetic variances were multiplied by a factor $Q(N_i^k/N_c)$, where

$$Q(x) = \begin{cases} 0 & \text{if } x < 0, \\ 10x^3 - 15x^4 + 6x^5 & \text{if } 0 \le x \le 1, \\ 1 & \text{if } x > 1. \end{cases} \tag{S53}$$

This is a smoothed step function: it is 0 for $x < 0$, increases monotonically to one at $x = 1$, and stays equal to 1 for $x > 1$ (Figure S2). It is constructed to be twice continuously differentiable for all $x$.

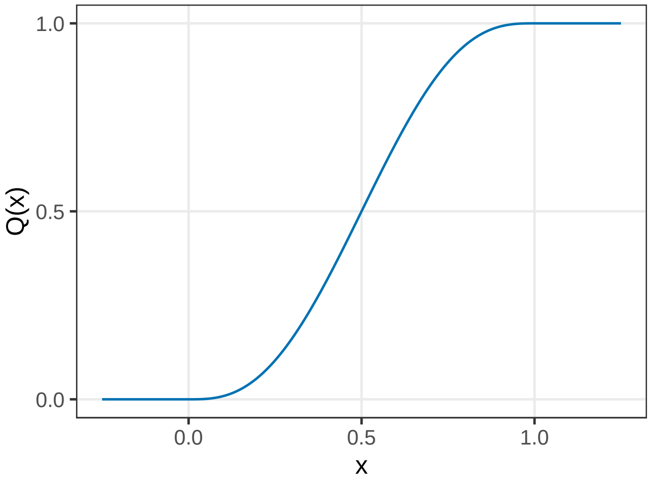


**Figure S2:** The smooth step function $Q(x)$, defined by Eq. S53. It is constructed to be piecewise polynomial and twice continuously differentiable everywhere.

This means that heritabilities were also modified below $N_c$, being instead given by the patch-dependent expression

$$h_i^2 = \frac{Q(N_i^k/N_c)V_{G,i}}{Q(N_i^k/N_c)V_{G,i} + V_{E,i}}. \tag{S54}$$

##### 3.5 Spatial structure and dispersal rates

We discretize latitudinal position into $L$ equidistant patches: patch 1 is at the pole, patch $L$ at the equator, with the rest linearly spaced in between. Migration happens between adjacent patches with a species-specific rate $d_i$:

$$m_i^{kl} = \begin{cases} d_i & \text{if } k = l \pm 1, \\ 0 & \text{otherwise.} \end{cases} \qquad (k, l = 1, \ldots, L) \tag{S55}$$

To the approximation that the two hemispheres of Earth are symmetric with respect to their temperature profiles, the pole-to-equator spatial profile could be repeated from equator to the other pole again (in reverse), and then wrapped around the whole planet. Due to this symmetry and periodicity, the range going from pole to equator already contains all information, and is the only part that needs to be modeled, provided one implements appropriate boundary conditions to account for the periodicity. Specifically, for patches 1 and $L$, the following replacements are made compared to Eqs. S36-S37:

$$\frac{\mathrm{d}N_i^1}{\mathrm{d}t} \to \frac{\mathrm{d}N_i^1}{\mathrm{d}t} + m_i^{1,2}N_i^2 - m_i^{2,1}N_i^1, \tag{S56}$$

$$\frac{\mathrm{d}N_i^L}{\mathrm{d}t} \to \frac{\mathrm{d}N_i^L}{\mathrm{d}t} + m_i^{L,L-1}N_i^{L-1} - m_i^{L-1,L}N_i^L, \tag{S57}$$

$$\frac{\mathrm{d}\mu_i^1}{\mathrm{d}t} \to \frac{\mathrm{d}\mu_i^1}{\mathrm{d}t} + h_i^2 m_i^{1,2}\frac{N_i^2}{N_i^1}(\mu_i^2 - \mu_i^1), \tag{S58}$$

$$\frac{\mathrm{d}\mu_i^L}{\mathrm{d}t} \to \frac{\mathrm{d}\mu_i^L}{\mathrm{d}t} + h_i^2 m_i^{L,L-1}\frac{N_i^{L-1}}{N_i^L}(\mu_i^{L-1} - \mu_i^L). \tag{S59}$$

##### 3.6 Climate model

Let $x$ denote latitudinal position, measured as the latitudinal distance from the north pole. The temperature at position $x$ and time $t$ is given by $T(x,t)$. The local temperature $T^k$ in patch $k$ is the one at the latitude corresponding to patch $k$. The components of the climate change model are as follows:

1. Temperature is initially constant in each patch, for a burn-in period of $t_0$ years. The initial average temperature $T_0(x)$ is $T_{\text{min}}$ at the pole, $T_{\text{max}}$ at the equator (Table S1), and linearly extrapolated in between:

$$T_0(x) = (T_{\text{max}} - T_{\text{min}})\frac{x}{X} + T_{\text{min}}, \tag{S60}$$

   where $X$ is the distance from pole to equator, measured in the same units as $x$.

2. Climate change starts after the burn-in period. In each patch, the qualitative shape of the relative temperature rise is given by the smoothed step function $Q((t-t_0)/t_E)$ (Eq. S53), where $t_0$ is the moment of the onset of climate change, and $t_E$ is the end of it.

3. Climate change stops after $t_E = 300$ years.

4. The extent of the temperature increase depends on latitude. This latitudinal trend is predicted with high confidence for the northern hemisphere for the next century, and is observed in all of IPCC's emission scenarios (IPCC 2013, chapter 12). We assume that the latitude-dependence of the final temperature increase (at $t_E$) is linear:

$$C_E(x) = (C_{\text{min}} - C_{\text{max}})\frac{x}{X} + C_{\text{max}}, \tag{S61}$$

   where $C_{\text{max}}$ is the maximum temperature increase (at the pole) and $C_{\text{min}}$ is the minimum increase (at the equator). The values of $C_{\text{max}}$ and $C_{\text{min}}$ (Table S1) are based on region-specific predictions of increase in temperature by 2100, in combination with estimates giving approximate increase by 2300, for the IPCC intermediate emission scenario (IPCC 2013, chapter 12).

Considering all the above, the time-dependence of the temperature profile $T(x,t)$ is written compactly as

$$T(x,t) = T_0(x) + C_E(x)Q\left(\frac{t-t_0}{t_E}\right). \tag{S62}$$

##### 3.7 Initial conditions

Both for resource species $1,2,\ldots,S$ and for consumer species $1,2,\ldots,S$, the initial temperature adaptations $\mu_i^k(0)$ are determined by the initial temperature profile such that species 1 is adapted to the polar temperatures, species $S$ to equatorial temperatures, and all other species are linearly spaced in between:

$$\mu_i^k(0) = T_0(i/S) = (T_{\text{max}} - T_{\text{min}})\frac{i}{S} + T_{\text{min}} \tag{S63}$$

at all spatial locations where the species is initially present. Initial population densities are centered at the location to which the species is best adapted, and are normally distributed around this point in space:

$$N_i^k(0) = \exp\left(-\frac{(\mu_i^k(0) - T_0(k/L))^2}{2\times 2^2}\right). \tag{S64}$$

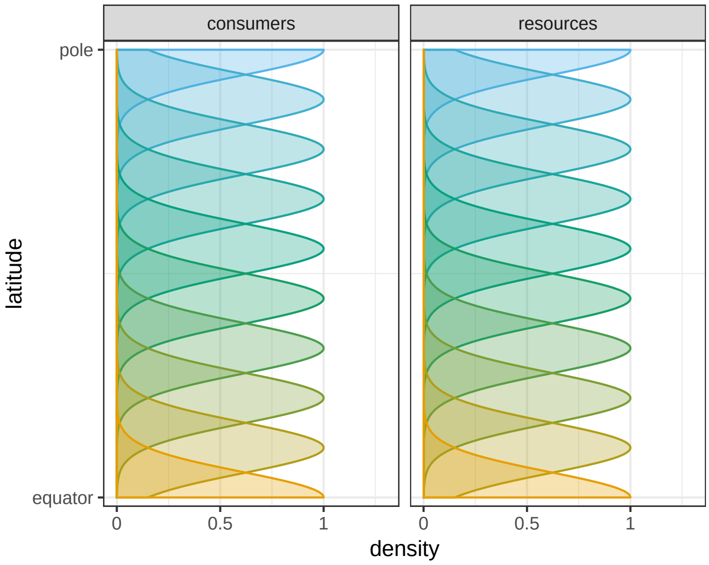


**Figure S3:** Initial density distribution of ten consumer (left) and ten resource species (right) along the latitudinal gradient (ordinate). The abscissa shows the population density at the corresponding latitude. Colors denote species, with cooler shades standing for more cold-adapted species.

### 4 Community response capacity

Let there be $S$ species distributed across $L$ patches, evolving according to our model. We denote the relative density of species $i$ in patch $k$ at time $t$ by $n_i^k(t)$:

$$n_i^k(t) = \frac{N_i^k(t)}{\sum_{j=1}^{S}\sum_{l=1}^{L} N_j^l(t)}. \tag{S65}$$

We then define the density-weighted trait lag, $\mathscr{A}^k(t)$, as

$$\mathscr{A}^k(t) = \frac{1}{S}\sum_{i=1}^{S} n_i^k(t)\left[\mu_i^k(t) - T^k(t)\right]^2, \tag{S66}$$

where $T^k(t)$ is the local temperature in patch $k$ at time $t$. This quantity measures the extent to which species match their environment, with more common species contributing more to the overall lag. $\mathscr{A}^k(t)$ may be further averaged across patches and time, yielding

$$\mathscr{A} = \frac{1}{L}\sum_{k=1}^{L}\frac{1}{t_E}\sum_{t=t_0}^{t_0+t_E}\mathscr{A}^k(t). \tag{S67}$$

The time averaging moves from the onset to the end of climate change, i.e., between years $t_0 = 1000$ and $t_0 + t_E = 1300$ (Table S1). In turn, we define the density-weighted dispersion of mean trait values, $\mathscr{V}^k(t)$, as

$$\mathscr{V}^k(t) = \frac{1}{S}\sum_{i=1}^{S} n_i^k(t)\left[\mu_i^k(t) - \bar{\mu}^k(t)\right]^2, \tag{S68}$$

where

$$\bar{\mu}^k(t) = \frac{1}{S}\sum_{i=1}^{S} n_i^k(t)\mu_i^k(t) \tag{S69}$$

is the weighted average of species' mean temperature optima in patch $k$. The more different species are in their trait means (with relatively more common species getting more weight for being different), the larger this value becomes. Again, we can average this quantity across time and patches to obtain

$$\mathscr{V} = \frac{1}{L}\sum_{k=1}^{L}\frac{1}{t_E}\sum_{t=t_0}^{t_0+t_E}\mathscr{V}^k(t). \tag{S70}$$

In Figure 6 of the main text, $\mathscr{A}$ is plotted against $\mathscr{V}$ to reveal that the more dispersed species' mean temperature optima are, the smaller the overall trait lag in a community. Communities with more varied mean temperature optima thus tend to be better adapted to local conditions. It also reveals that certain model types (in particular, models without temperature-dependent competition) consistently produce larger mean trait dispersion $\mathscr{V}$ than others. Moreover, this relationship is independent of species' average dispersal abilities and genetic variances: when varying these parameters in a much wider range than in the main text (Figure S4), the same regression line is retained. A similar linear relationship as the main text's Figure 6 holds for the consumer species as well (Figure S5), though with a different slope and intercept.

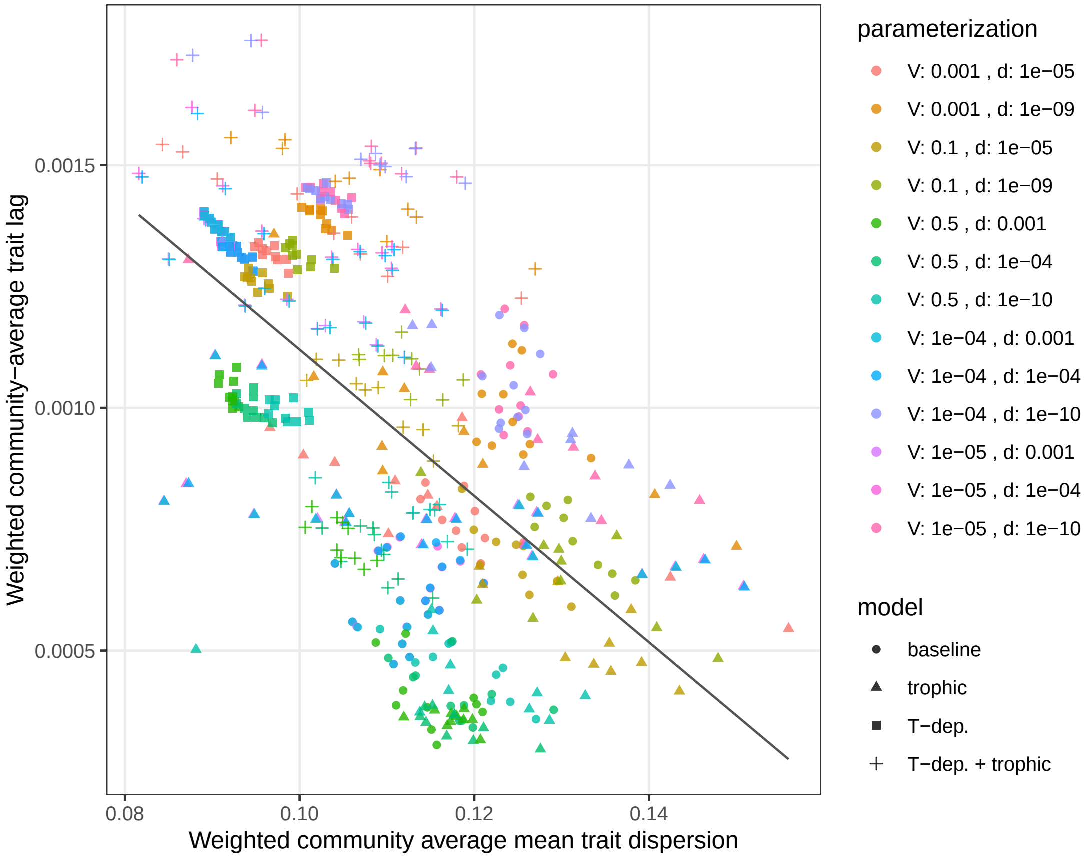


**Figure S4:** As Figure 6 in the main text, but for a much wider range of dispersal abilities and genetic variances. Even so, the regression line (slope $-0.015$, intercept 0.0026, $p < 2.210^{-16}$) is the same as before, indicating that these two parameters leave the relationship unaffected.

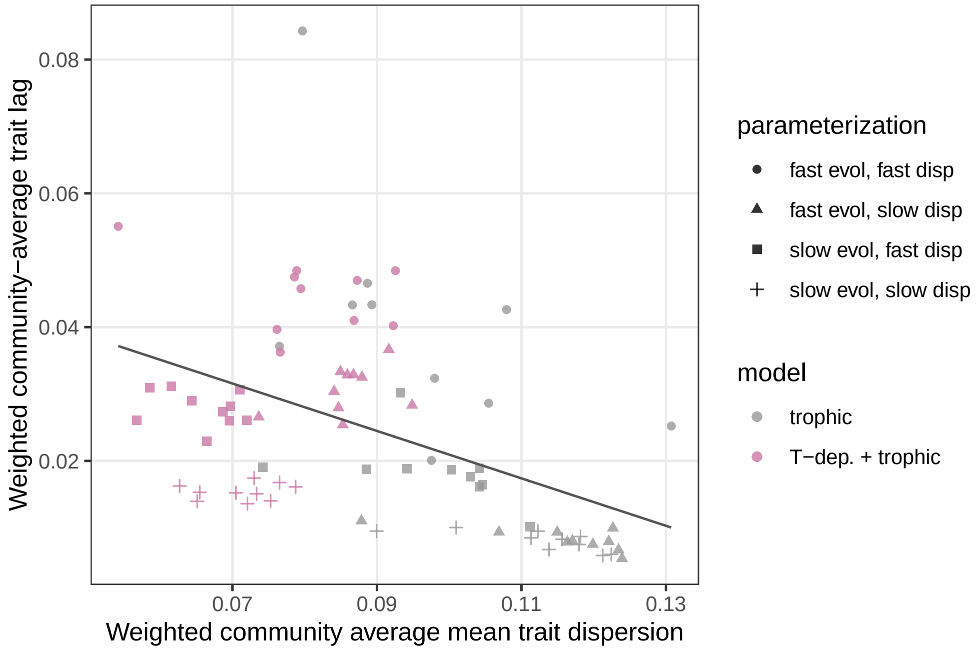


**Figure S5:** As Figure 6 in the main text, except for consumer species.

### 5 Additional results

#### 5.1 Community turnover

Species turnover in time is measured by the Jaccard distance. Let $\mathscr{S}(t)$ denote the set of species present at time $t$, and $|\mathscr{S}(t)|$ the number of elements in the set. The Jaccard distance $J(t_0,t)$ between the community state at some reference time $t_0$ and a later time $t$ reads

$$J(t_0,t) = 1 - \frac{|\mathscr{S}(t_0) \cap \mathscr{S}(t)|}{|\mathscr{S}(t_0) \cup \mathscr{S}(t)|}, \tag{S71}$$

where $\cap$ is the intersection and $\cup$ the union of two sets. This formula is agnostic about the spatial extent of a community: it can be applied to each patch separately, or the community as a whole. An in-between approach is to obtain regional turnover by applying Eq. S71 to the top third of all $L$ patches (the polar region), the middle third (temperate region), and bottom third (tropical region). Figure S6 shows how regional species compositions change in time, with the community state at the start of climate change playing the role of the reference community at $t_0$.

#### 5.2 Summary plots for consumers

Figures S7-S9 are like Figures 3-5 in the main text, except they show the distribution ranges, local diversity, and global diversity information for the consumer instead of the resource species. Figure S10 is the analogue of Figure S6 for consumers.

#### 5.3 Results with 30 species per trophic level

Figures 3-6 in the main text assume 50 species per trophic level. This number is sufficient for all relevant community patterns to stabilize, such that adding more species has no more substantial effect. To show this, we display species ranges, local richness, and global richness for $S = 30$ species per trophic level (Figures S11-S20). The qualitative patterns are identical to those in Figures 3-6 of the main text, and Figures S6-S10.

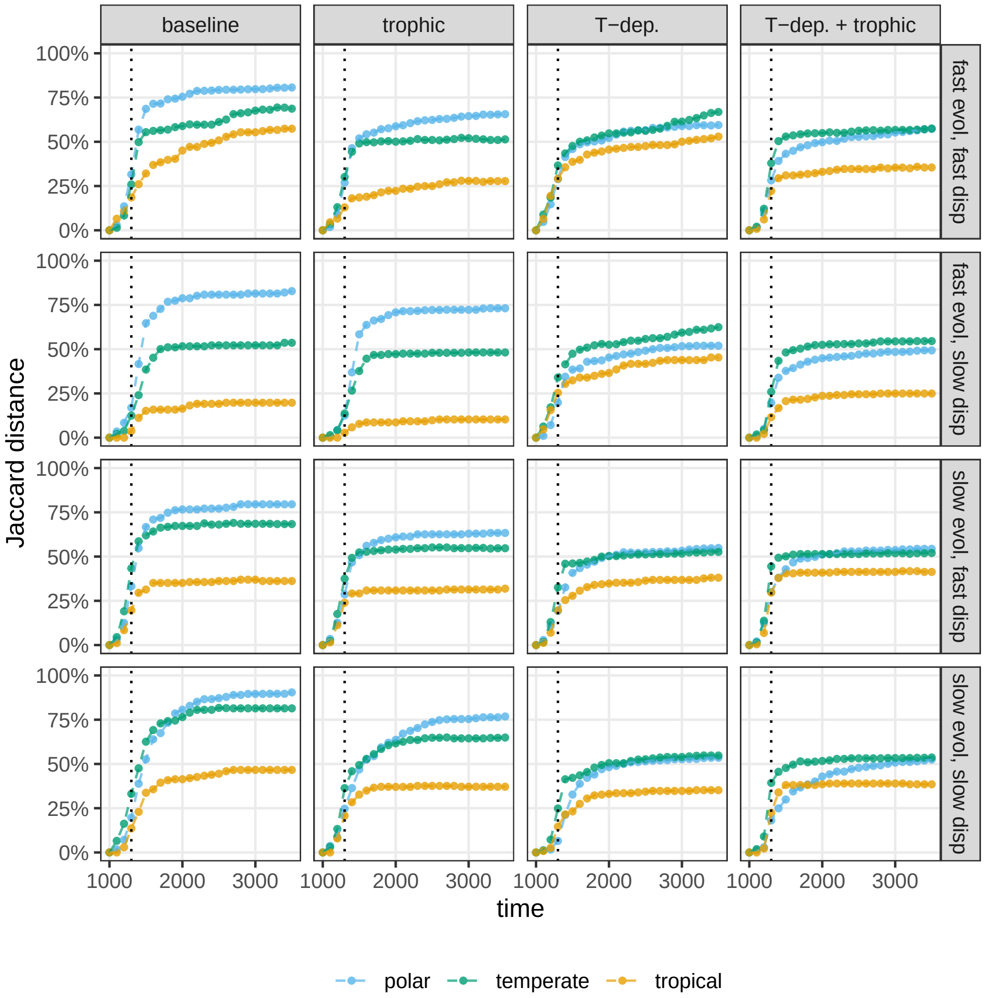


**Figure S6:** Jaccard distance between the regional community composition of resource species at the start of climate change and subsequent times, in steps of 100 years (points; dashed lines connect them for better visibility). A value of 0% means that the same set of species is present as at the start, while 100% indicates a complete turnover in regional species composition. The Jaccard distance is averaged over patches (merged into regions, indicated by color) and replicates. Columns show different ecological models, rows various parameter combinations of average dispersal ability and available genetic variance.

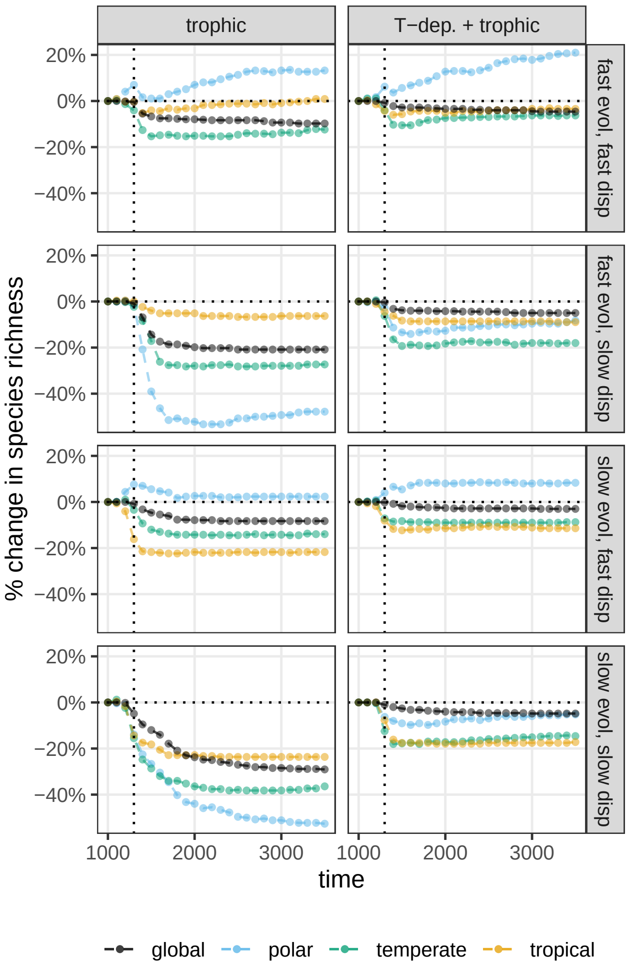


**Figure S7:** As Figure 3 of the main text, but for consumer species (species 51 to 100).

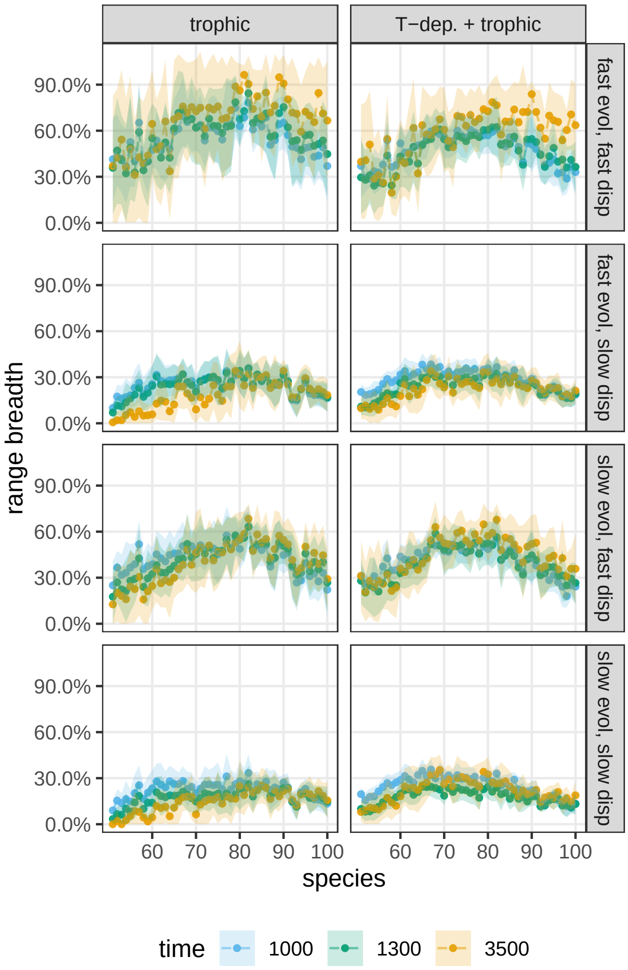


**Figure S8:** As Figure 4 of the main text, but for consumer species (species 51 to 100).

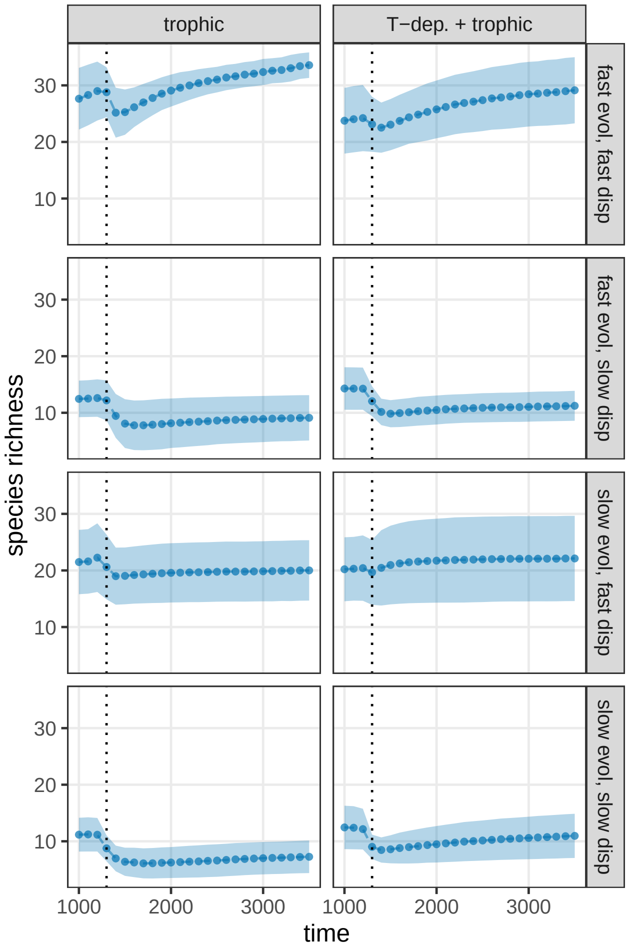


**Figure S9:** As Figure 5 of the main text, but for consumer species (species 51 to 100).

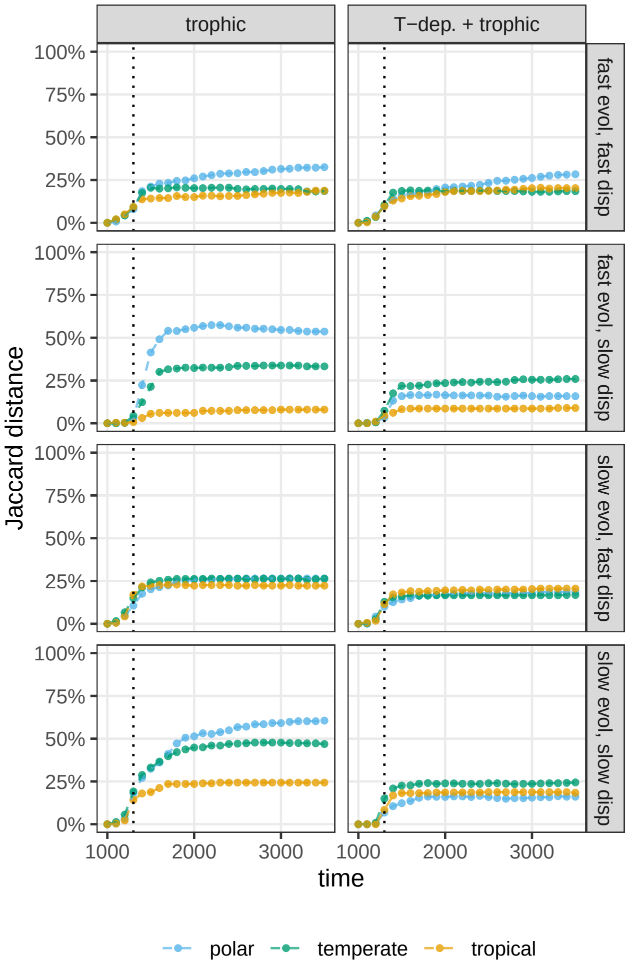


**Figure S10:** As Figure S6, but for consumer species (species 51 to 100).

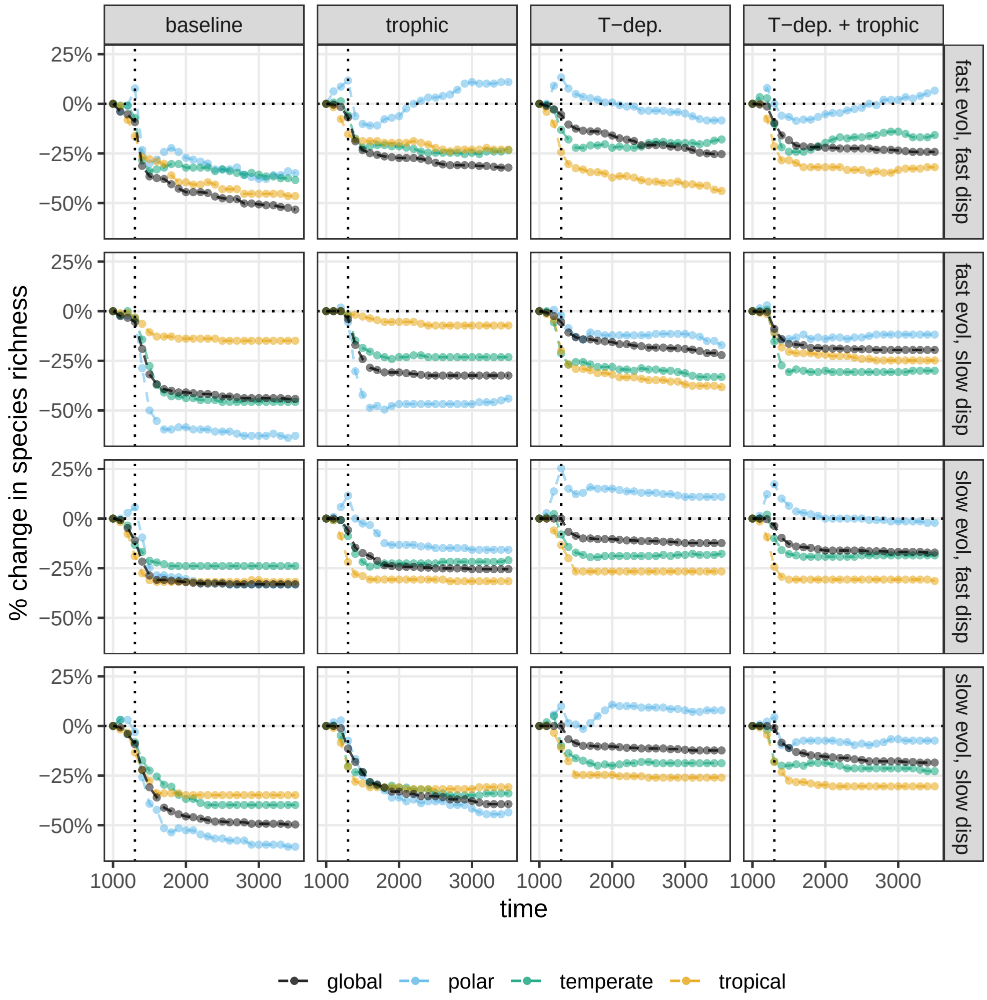


**Figure S11:** As Figure 3 of the main text, but with $S = 30$ resource species.

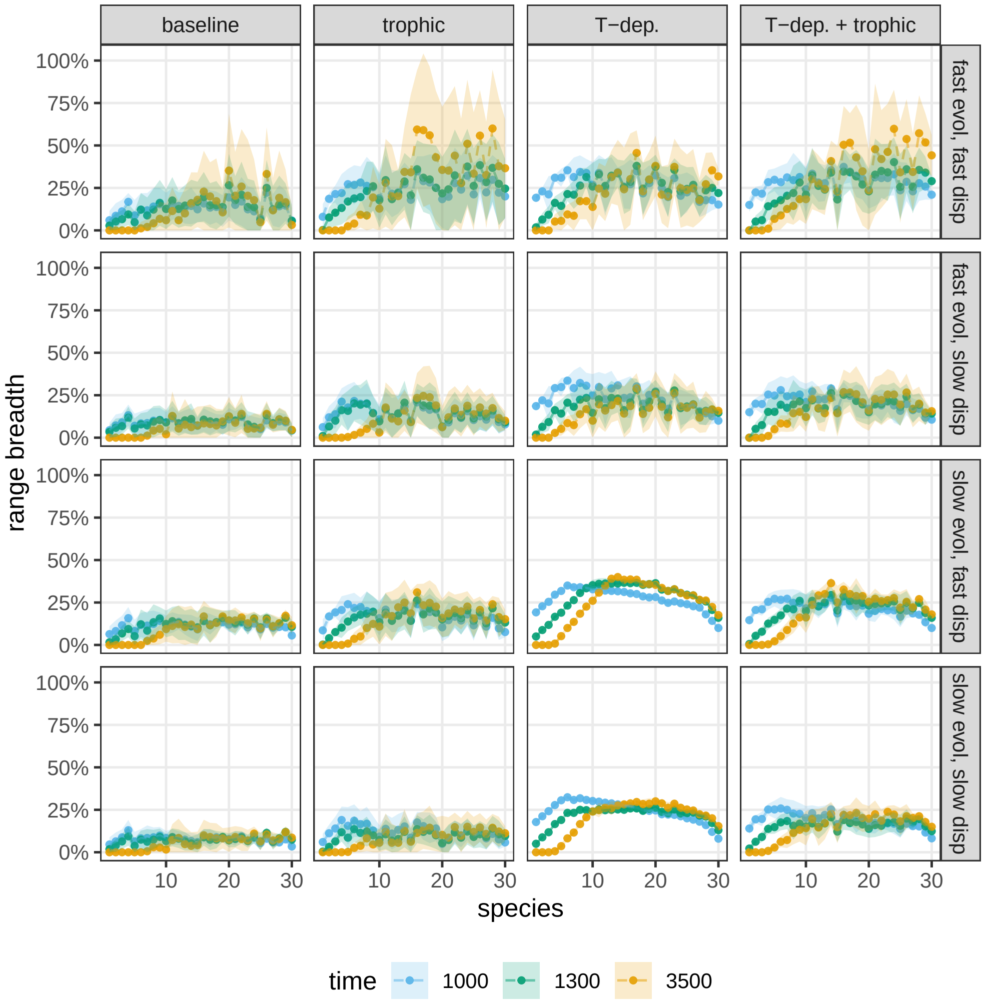


**Figure S12:** As Figure 4 of the main text, but with $S = 30$ resource species.

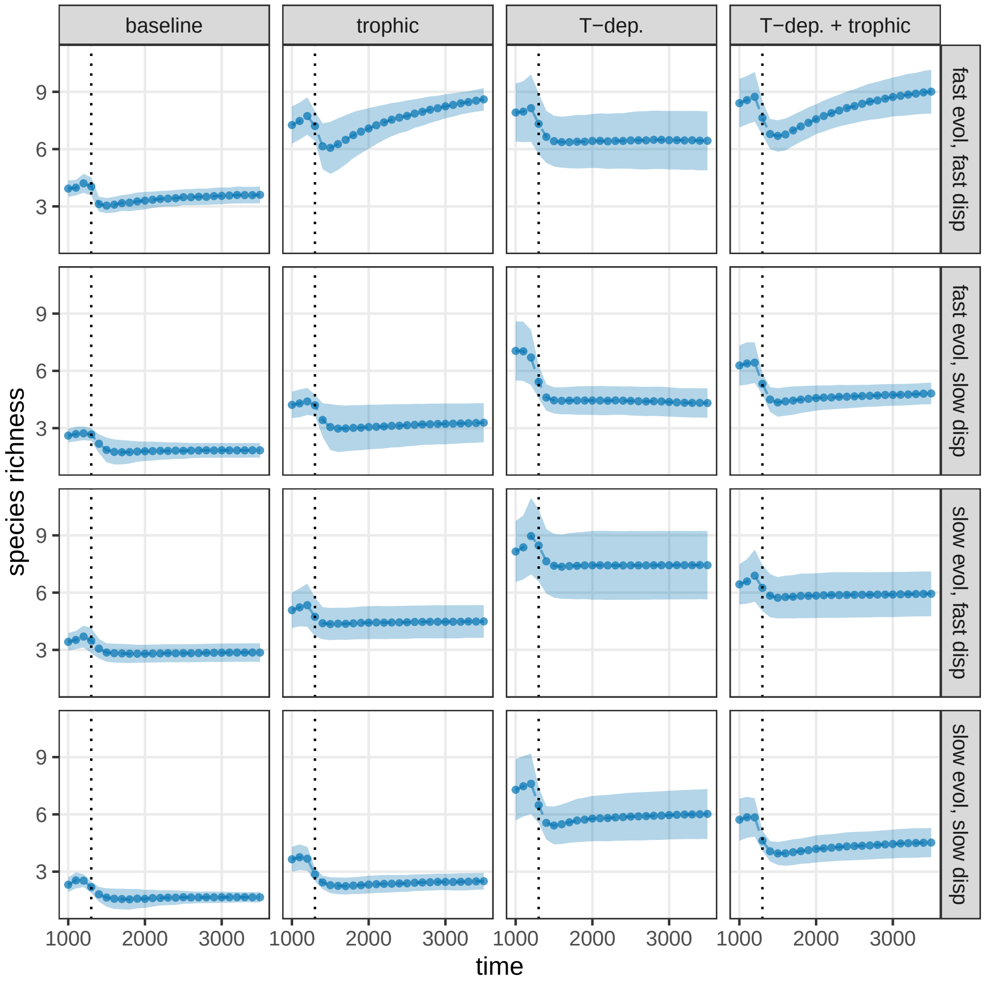


**Figure S13:** As Figure 5 of the main text, but with $S = 30$ resource species.

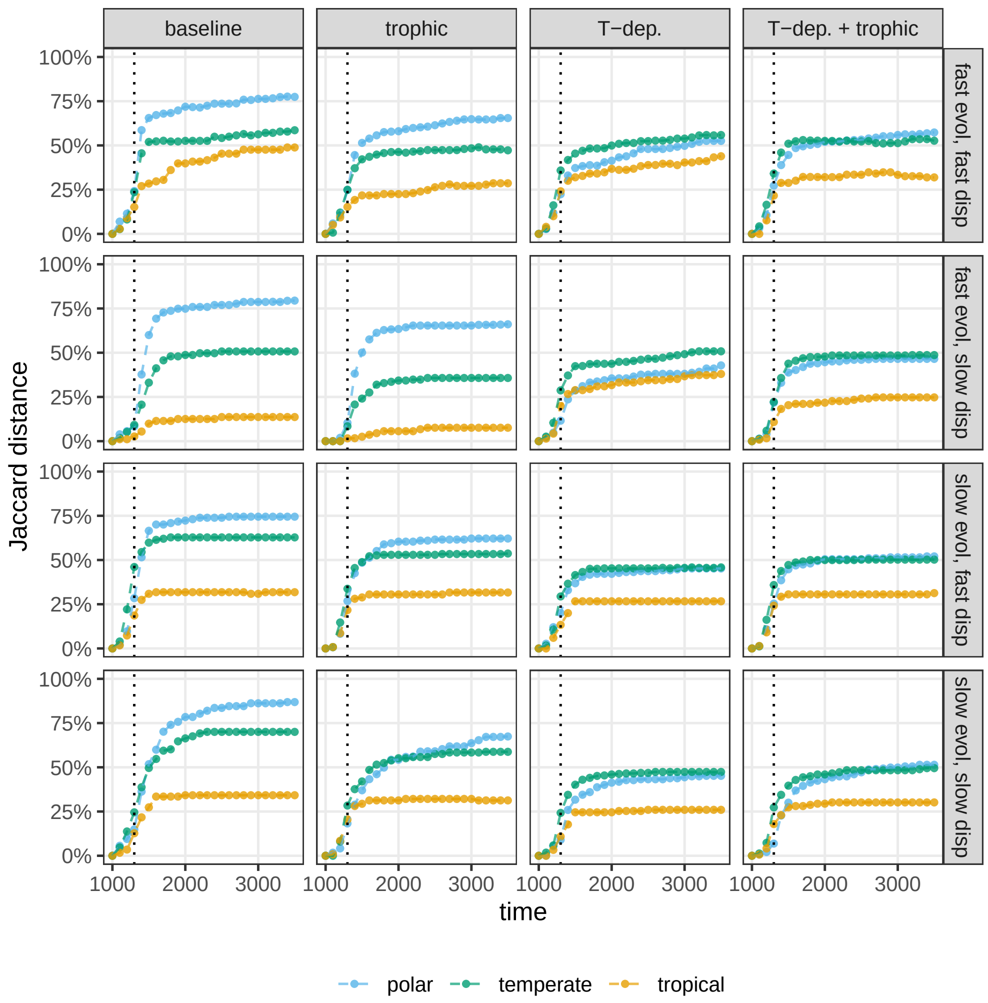


**Figure S14:** As Figure S6, but with $S = 30$ resource species.

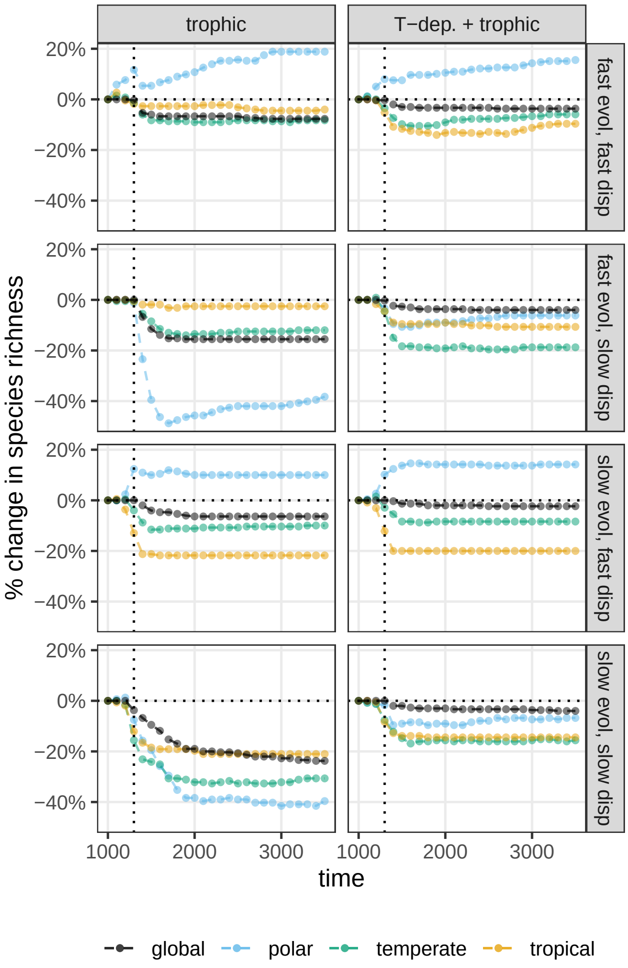


**Figure S15:** As Figure S7, but with $S = 30$ consumer species (species 31 to 60).

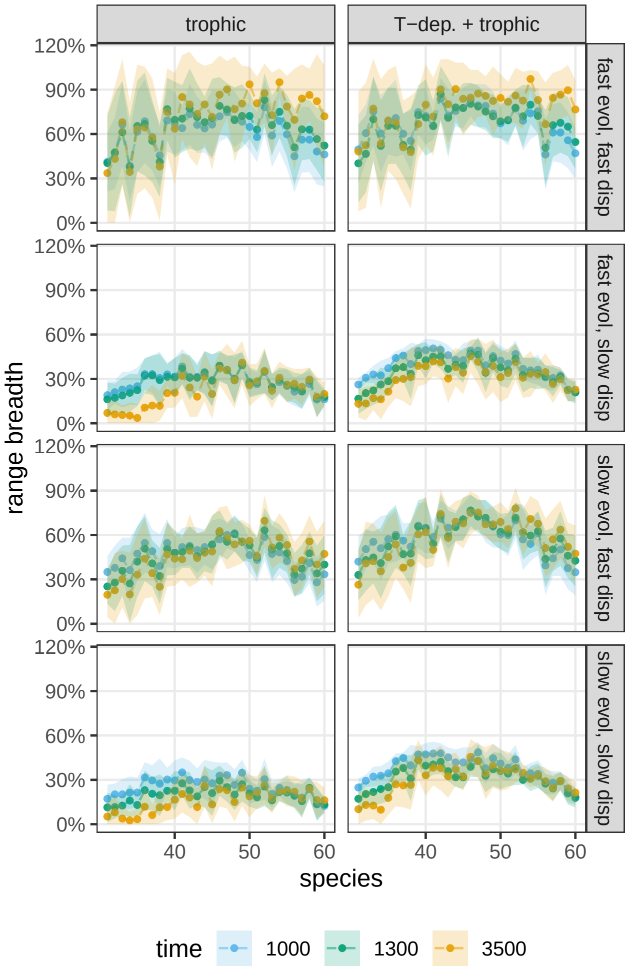


**Figure S16:** As Figure S8, but with $S = 30$ consumer species (species 31 to 60).

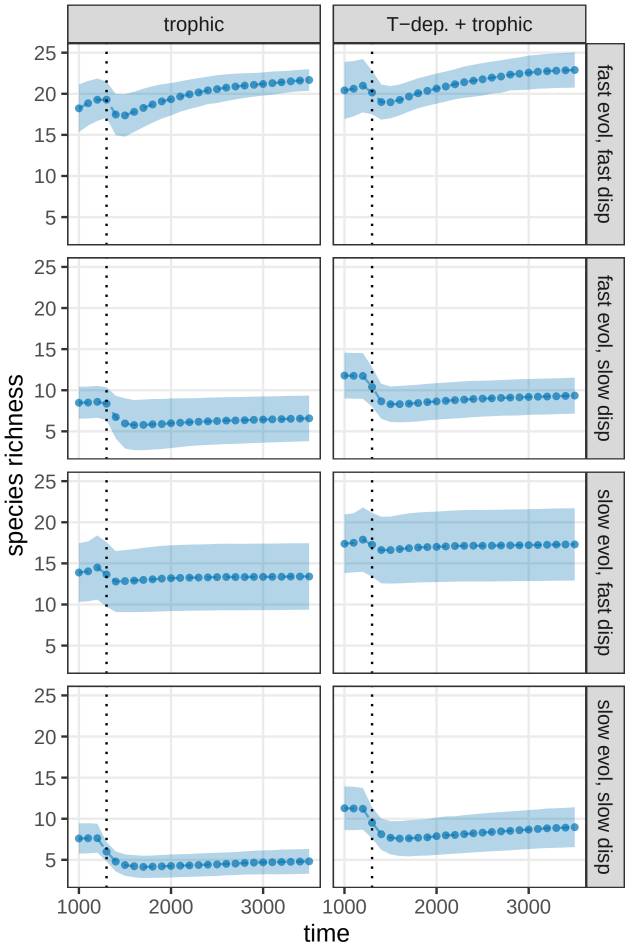


**Figure S17:** As Figure S9, but with $S = 30$ consumer species (species 31 to 60).

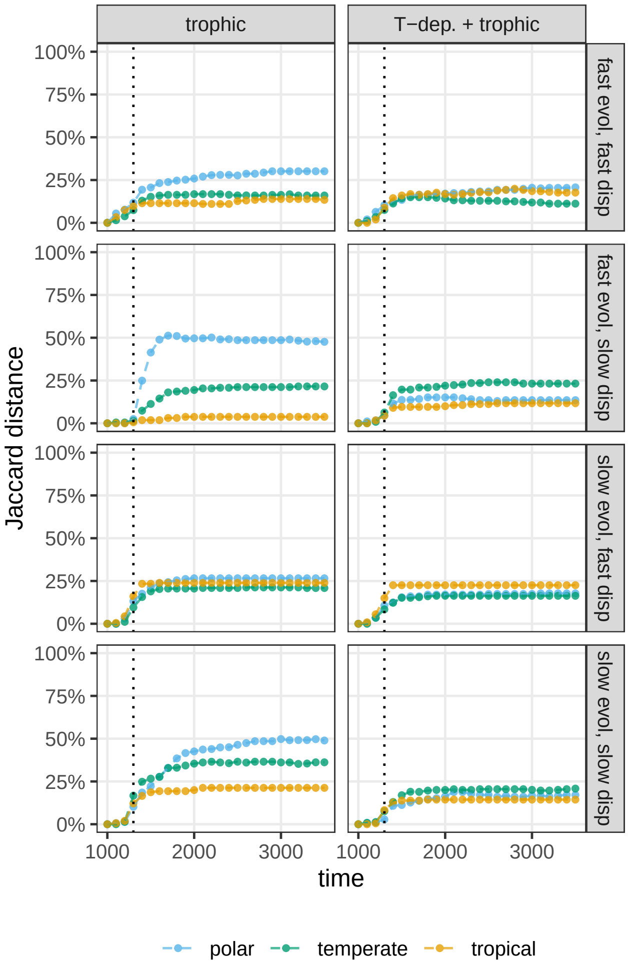


**Figure S18:** As Figure S10, but with $S = 30$ consumer species (species 31 to 60).

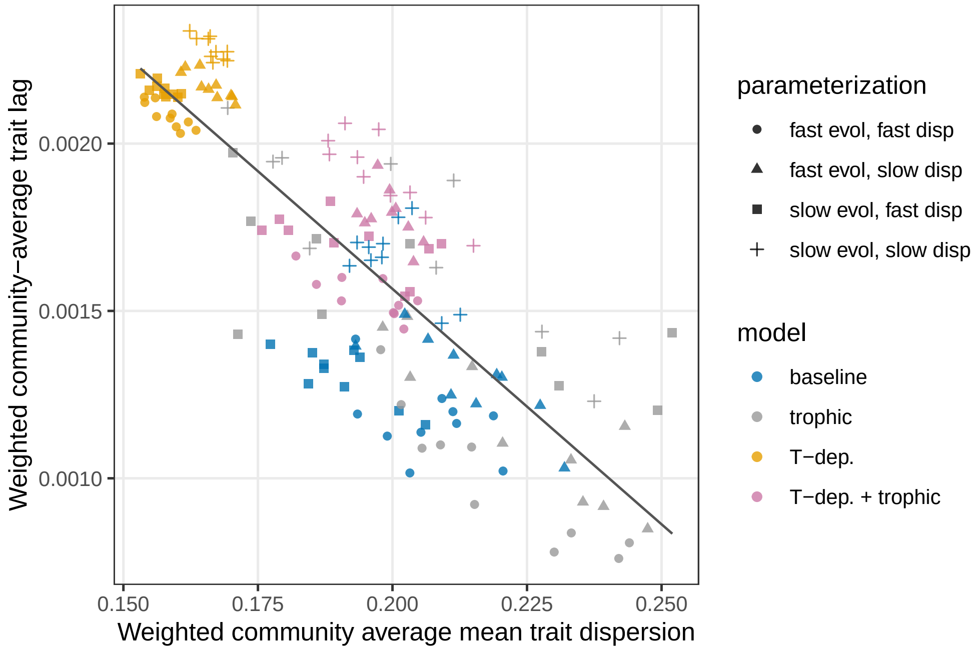


**Figure S19:** As Figure 6 in the main text, but with $S = 30$ resource species.

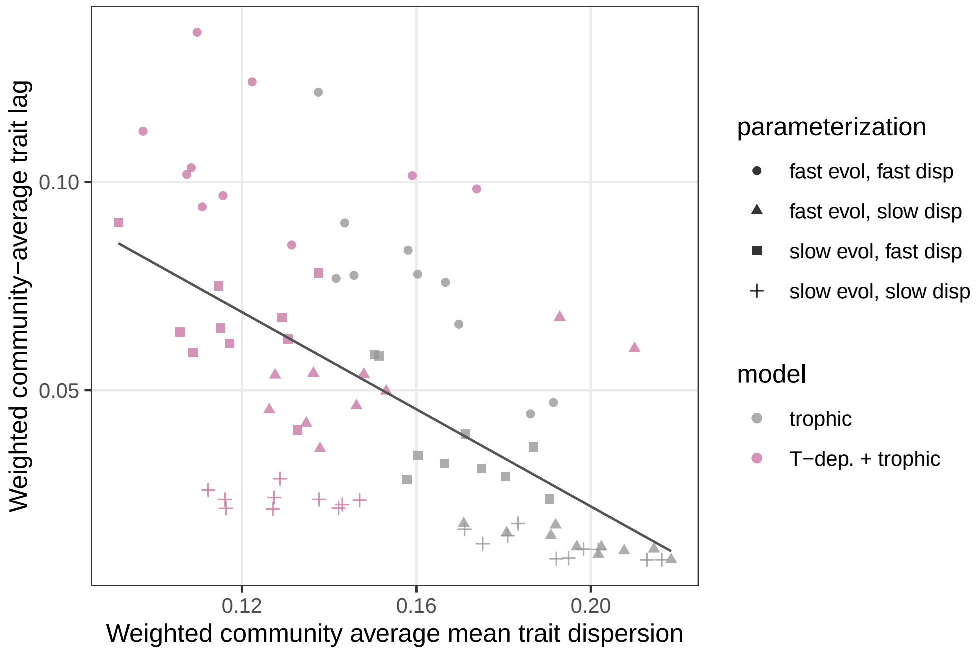


**Figure S20:** As Figure S5, but with $S = 30$ consumer species (species 31 to 60).

### 6 Parameter values

| parameter | value | units | comment |
|---|---|---|---|
| $S$ | 50 / 50 | | no. of species (resource / consumer) |
| $L$ | 50 | | no. of patches along latitudinal gradient |
| $X$ | $10^7$ | m | distance from pole to equator |
| $\rho_i$ | [0.9, 1.1] / [0.09, 0.11] | °C yr$^{-1}$ | tradeoff parameter (resources / consumers) |
| $b_w$ | 4.0 | °C | tolerance width's trait-dependence intercept |
| $a_w$ | 0.1 | | tolerance width's trait-dependence slope |
| $\kappa_i$ | 0.1 | yr$^{-1}$ | intrinsic mortality |
| $\eta$ | 1 | °C | competition width |
| $\varepsilon_i$ | 0.05 | | consumer conversion efficiency |
| $q_i$ | [1, 10] | g$^{-1}$yr$^{-1}$ | attack rate |
| $H_i$ | [0.5, 1] | yr | handling time |
| $W_{ij}$ | 1 if $i$ eats $j$; else 0 | | adjacency matrix of feeding network |
| $\omega_{ij}$ | [# of $i$'s resources]$^{-1}$ | | relative consumption rate of $i$ on $j$ |
| $\overline{V}$ | $10^{-1}$ or $10^{-3}$ | °C$^2$ | mean genetic variance |
| $\overline{d}$ | 100 or 0.01 | m yr$^{-1}$ | mean annual dispersal distance |
| $a_{ii}$ | [0.1, 0.3] | g$^{-1}$yr$^{-1}$ | intraspecific comp. coeffs. (resources only) |
| $a_{ij,\, j \neq i}$ | [0.075, 0.225] | g$^{-1}$yr$^{-1}$ | interspecific comp. coeffs. ($i$, $j$ resources) |
| $N_c$ | $10^{-5}$ | | threshold density for heritability reduction |
| $T_{\min}$ | $-10$ | °C | initial mean temperature at pole |
| $T_{\max}$ | 25 | °C | initial mean temperature at equator |
| $C_{\min}$ | 1.26 | °C | projected temperature increase at pole |
| $C_{\max}$ | 9.66 | °C | projected temperature increase at equator |
| $t_0$ | 1000 | yr | time at which climate change starts |
| $t_E$ | 300 | yr | end of climate change (after $t_0$) |

**Table S1:** Parameter values. Unit abbreviations: g (grams), m (meters), °C (Celsius degrees), yr (years). When intervals are shown, values are uniformly drawn for each species.

| parameter | value | units | comment |
|---|---|---|---|
| $V_{G,i}$ | $[0.5\overline{V},\, 1.5\overline{V}]$ | °C$^2$ | genetic variance |
| $V_{E,i}$ | $\overline{V}$ | °C$^2$ | environmental variance |
| $\sigma_i^2$ | $V_{G,i} + V_{E,i}$ | °C$^2$ | total phenotypic variance |
| $h_i^2$ | $V_{G,i}/\sigma_i^2$ | | heritability |
| $d_i$ | $[0.1\overline{d},\, 10\overline{d}]$ | m yr$^{-1}$ | annual dispersal distance |
| $m_i^{kl}$ | $\begin{cases} d_i & \text{if } k = l \pm 1 \\ 0 & \text{otherwise} \end{cases}$ | m yr$^{-1}$ | dispersal matrix (for $k, l$ in $1, 2, \ldots, L$) |

**Table S2:** Derived parameters, using the values in Table S1. Notation and unit abbreviations are as in Table S1.
